# Spatial and temporal constraints on the composition of microbial communities in subsurface boreholes of the Edgar Experimental Mine

**DOI:** 10.1101/2021.06.09.447828

**Authors:** Patrick H. Thieringer, Alexander S. Honeyman, John R. Spear

**Affiliations:** Department of Civil and Environmental Engineering, Colorado School of Mines, Golden, Colorado, USA

**Author notes:** Adress correspondence to John R. Spear.

## Abstract

The deep biosphere hosts uniquely adapted microorganisms overcoming geochemical extremes at significant depths within the crust of the Earth. While numerous novel microbial members with unique physiological modifications remain to be identified, even greater attention is required to understand the near-subsurface and its continuity with surface systems. This raises key questions about networking of surface hydrology, geochemistry affecting near-subsurface microbial composition, and resiliency of subsurface ecosystems. Here, we apply molecular and geochemical approaches to determine temporal microbial composition and environmental conditions of filtered borehole fluid from the Edgar Experimental Mine (∼150 meters below the surface) in Idaho Springs, CO. Samples were collected over a 4-year collection period from expandable packers deployed to accumulate fluid in previously drilled boreholes located centimeters to meters apart, revealing temporal evolution of borehole microbiology. Meteoric water feeding boreholes demonstrated variable recharge rates likely due to a complex and undefined fracture system within the host rock. 16S rRNA gene analysis determined unique microbial communities occupy the four boreholes examined. Two boreholes yielded sequences revealing the presence of *Proteobacteria, Firmicutes,* and *Nanoarcheota* associated with endemic subsurface communities. Two other boreholes presented sequences related to soil-originating microbiota, which likely indicate a direct link to surface infiltration. High concentrations of sulfate suggest sulfur-related metabolic strategies dominate within these near-subsurface boreholes. Overall, results indicate microbial community composition in the near-subsurface is highly dynamic at very fine spatial scales (<20cm) within fluid-rock equilibrated boreholes, which additionally supports the role of a relationship for surface geochemical processes infiltrating and influencing subsurface environments.

**Importance:** The Edgar Experimental Mine, Idaho Springs, Colorado provides inexpensive and open access to borehole investigations for subsurface microbiology studies. Understanding how microbial processes in the near-subsurface are connected to surface hydrological influences like meteoric input is lacking. Investigating microbial communities of subsurface mine boreholes provides evidence of how geochemical processes are linked to biogeochemical processes within each borehole, and the geochemical connectedness and mobility of surface influences. This study details microbial community composition and fluid geochemistry over spatial and temporal scales from boreholes within the Edgar Mine. These findings are relevant to biogeochemistry of near-surface mines, caves and other voids across planetary terrestrial systems. In addition, this work can lead to understanding how microbial communities relating to both fluid-rock equilibration and geochemical influences may enhance our understanding of subsurface molecular biological tools that aid mining economic practices to reflect biological signals for lucrative veins in the near subsurface.

## Introduction

Fervor to constrain and discover microbial habitability in the subsurface outweighs our understanding of major biogeochemical drivers supporting growth for microorganisms in the subsurface. Field sites spreading terrestrial, lacustrine, hot springs, and marine environments have gained traction for molecular biological investigation due to unique, extreme conditions mimicking potential environmental settings of early life on Earth as well as analogous to those on other planetary systems [1–4]. Locations such as the Lost City Hydrothermal Field have elucidated the interconnection between geological, chemical, and biological activity of marine subsurface microbial communities [5]. Marine research has documented microbial growth reaching depths up to 2 km below the seafloor, pushing previous expectations for the limits of microbial growth [6]. While marine systems have unveiled the extent of microbial subsurface habitability in oceanic settings, research aimed at understanding accessible terrestrial environments has gained considerable interest because of the more uniquely limiting constraints (low nutrient access, low O_2_ availability, and elevated temperatures) towards microbiological habitability.

The large fount of microbial life in the deep biosphere remains to be explored through terrestrial subsurface investigations [7]. Recent studies have constrained microbial biomass existing in the deep biosphere by quantifying deep fracture fluids from the Canadian Shield [8]. Terrestrial sites have increased our knowledge of where microbial ecosystems flourish, expanding to subsurface ultra-basic and high temperature serpentinized systems [9,10,11]. The terrestrial subsurface can offer more geologic and hydrologic variability compared to marine environments due to the presence of unconstrained fluid sourcing and lithology in surface soils and bedrock [7, 12–14]. Even more, water-rock interactions demonstrated at hydrothermal sites distinguish the chemical disparity between different subsurface exchanges that allow certain environments to become more suitable for hosting microorganisms [15]. These sites demonstrate chemolithotrophic processes that sustain ecosystems of low biomass to utilize unique sources of energy [16]. The contemporary outlook of an “extreme” environment encompasses some variation of a chemical or hydrological endpoint for microorganisms to exist primarily due to physiological constraints. However, new insights into subsurface and deep subsurface research may not necessitate harsh geochemical gradients or temperatures. Instead, the restricted access to nutrients, vital for microbial activity, makes these terrestrial subsurface studies of interest to investigate how moderately warm and/or acidic environments can still be uniquely constraining to microbiota.

Continental subsurface studies are beginning to unfold a considerable amount of evidence for understanding microbiology within deep fractures and aquifers [12]. However, the scope of which microorganisms survive in subsurface environments from meteoric water infiltration to introduce nutrients from the surface versus chemical energy sources originating in local settings remains insufficiently answered [17]. A common theme for subsurface exploration and microbial habitability continues to be the necessity for water. Water plays a vital role in the presence of microorganisms at any depth within the subsurface. When considering planetary studies in the search for life, water is a common choice as “follow the water” for high priority targets for investigation including the icy moons Europa and Enceladus [2].

In order to study subsurface microbiology, many deep subsurface observatories have been established. Reaching depths of up to ∼2.4 km, the Sanford Underground Research Laboratory in South Dakota has investigated the geochemistry and DNA sequences of up to 10,000 year-old circuitous groundwater to understand the energetics of chemolithotrophic organisms [15]. These subsurface laboratories extend across multiple continents and countries from the U.S., Finland, France, Japan, and the U.K. [18]. Another exists in Canada at the Kidd Creek Observatory, where fracture waters at 2.4 km depth were acquired in order to cultivate microorganisms from the fluids and determine the metabolic energetic pathways from parallel geochemical analysis [19]. Onstott et al. [17, 20] have established the presence of a microbial community in a gold mine in the deep subsurface from fluids obtained up to 3 km below the surface. These fluids proved microbiota indigenous to the host rock environment and further demonstrate the potential for life driven by equilibration of water-rock interactions.

The established observatories in the deep subsurface offer novel investigations into niche environmental conditions; however, they only shed light on chemical energy stored within local settings. Studies have utilized deep fracture fluids or ancient aquifers to constrain life in the subsurface, yet only limited research has considered the influence of meteoric water penetration into the near-subsurface. The role of meteoric water infiltration as a transportation mechanism for nutrients introduces questions regarding the greater potential for life, the connectivity of the surface to the subsurface and the relation of surface processes to subsurface microorganisms. Meteoric infiltration also offers the opportunity for recharge events, increasing the prospect for sampling and investigating controlling geochemical variables on subsurface habitability.

An additional consideration for deep observatories is the cost and accessibility of boreholes for sampling and inquiry. Due to expenses from boreholes and gaining access to subsurface environments, numerous questions remain unanswered from both marine and terrestrial subsurface ecosystem exploration [7]. Therefore, easier access into the subsurface is advantageous not only to establish consistent research regarding the microbial communities existing in the subsurface, but also to monitor geochemical variables influencing subsurface habitability with greater control over experimental design and statistical power.

This paper discusses the advantages of studying subsurface microbiology at the Edgar Experimental Mine, an easily accessible portal to the subsurface with ample opportunity to unveil and experiment with the subsurface biosphere. Meteoric water infiltrating the fractures of the mine leak out of the boreholes, offering the opportunity to study the subsurface microbiology of meteoric water influenced by fluid-rock equilibration (**Figure 1**). Additionally, the recharge of water from the surface makes it possible to study temporal influences through incubation and recharge periods. Here, we apply genomic, geochemical, and isotopic approaches to better define the variable microbial composition of borehole fluids and their relationship to surface influences from meteoric penetration into the subsurface.

**Figure 1.**
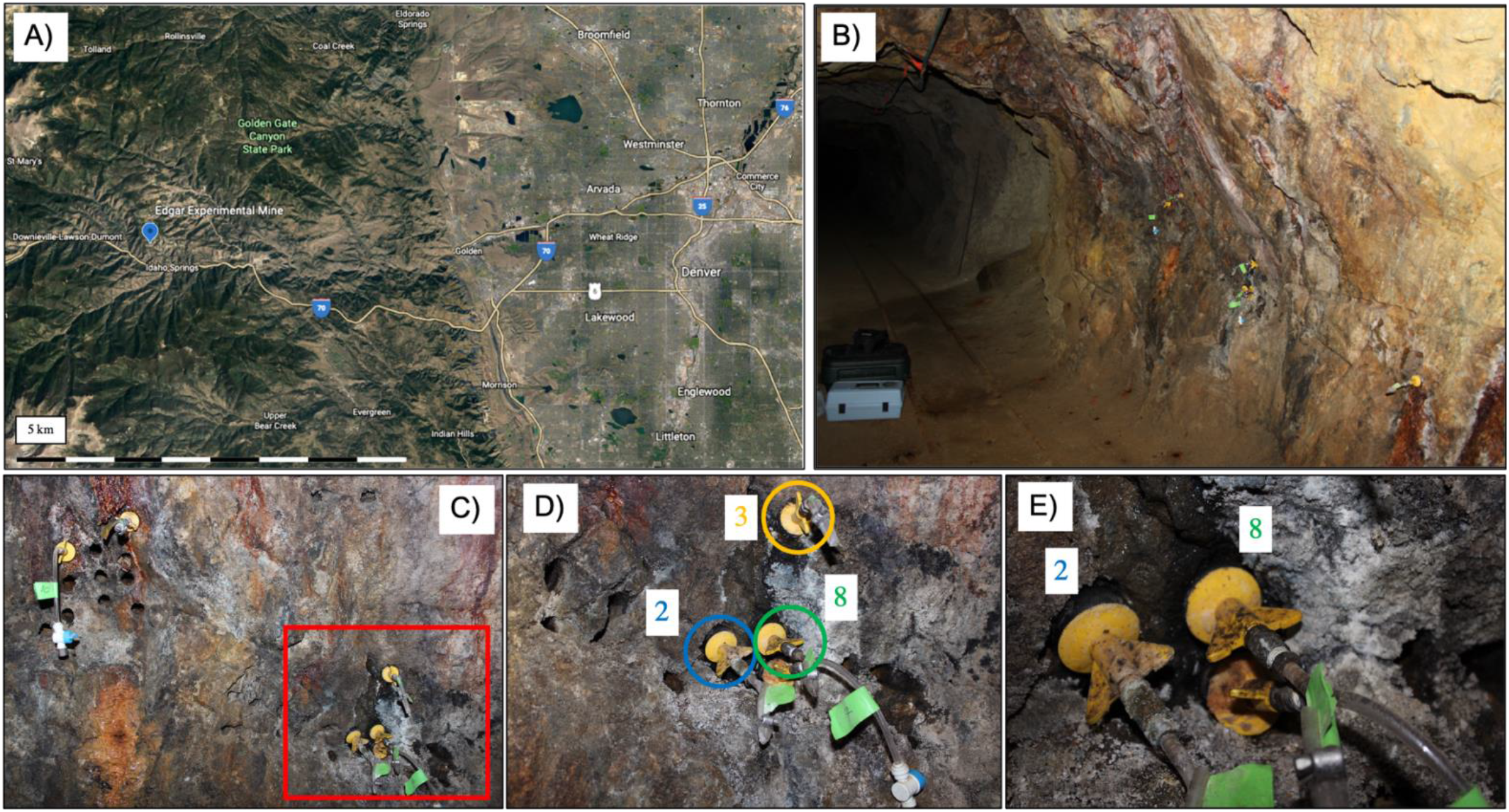
Composite images depicting the location of Edgar Experimental Mine and the boreholes sampled within this study. A) Landsat image of the location of Denver in reference to the mine in Idaho Springs following along I-70. B) The sampling site for the study located in the area termed “C-Right” within the mine. Packed boreholes can be seen extruding out of the wall where suspected fluid leaks out and can be plugged up in order to collect fluid for subsequent sampling events. C) Boreholes with packers included in the red box are boreholes observed in this study. Additional packed boreholes did not collect any fluid for sampling events. Previously drilled boreholes can be observed that do not leak any fluid potentially due to the complexity of the undefined fracture system. D) A close-up image of the red box indicated from the previous figure with boreholes 2, 3, and 8 observed in this study. Borehole 6 (not captured) is approximately 5 meters away on another wall of the field site. E) A closer image of boreholes 2 and 8 located centimeters apart from each other; however, each collects distinctly different amounts of fluid at varying rates of recharge.

## Results

### Borehole Geochemistry

Over the four-year sample collection period, the subsurface boreholes remained relatively uniform in their recorded analyte and physical chemistry measurements. While boreholes occasionally fluctuate in minor analyte concentrations, no discrete patterns were observed. Despite a lack of change, each borehole hosts nuanced geochemical profiles providing niche microenvironments for microbial communities. Fluid recharge in the boreholes was unpredictable regarding volume and temporal repetition. Borehole 2 was able to consistently fill with greater than 1 L of water each sampling period, whereas all other boreholes spanned months or years before refilling with fluid (**Figure 1**). The remaining boreholes did not refill with predictable volumes of water at each sampling period, further emphasizing a complex fracture pathway for fluid transport. Physical chemistry composition remained relatively homogenous amongst boreholes. Temperature ranges can be attributed to seasonal fluctuations, generally remaining between 10.6 to 13.5°C and reaching 17.6°C in summer. Dissolved oxygen varied most strongly from 1.97 to 7.7 mg/L, while pH was generally uniform within individual boreholes but varied amongst all samples ranging 5.30 to 7.16 pH.

Ion chromatography (IC) and inductively-coupled plasma atomic emission spectroscopy (ICP-AES) results of borehole fluids demonstrated weakly variable subsurface fluid chemistry. Concentrations of IC results (anions; putative electron acceptors for microbial growth as well as phosphorous sourcing) tend to be very low or undetected, allowing for niche substrate availability of microbial communities. F^−^ levels across all boreholes ranged from below detection or from 0.15 to 0.33 ppm. NO_2_ levels were undetected except for three samples that recorded values of 0.19, 0.27, and 0.37 ppm. NO_3_ remained undetected or reported values ranging from 0.45 to 1.27 ppm. Cl^−^ levels varied unexpectedly within individual boreholes with concentrations between 1.59 to 22.08 ppm, but spiked as high as 474.85 ppm. SO_4_ concentrations were the largest recorded analyte amongst all boreholes and consistently remained in high concentrations usually greater than 1000 ppm. Analytes chosen from ICP analysis were selected to demonstrate greatest variability amongst the borehole fluids (**Table 1**). In particular, the metals Mn and Zn were in the highest notable concentrations with maximum values of 27.15 mg/L and 18.46 mg/L respectively. There is a linear relationship between the dissolved Mn and Zn concentrations in the borehole fluids (**Figure 2)**. This highlights the distinction between the metal-rich fluids found in Boreholes 2 and 3 and more dilute fluids demonstrated in Boreholes 6 and 8. All recorded field measurements, IC, and ICP data are presented in **Table 1**.

**Figure 2.**
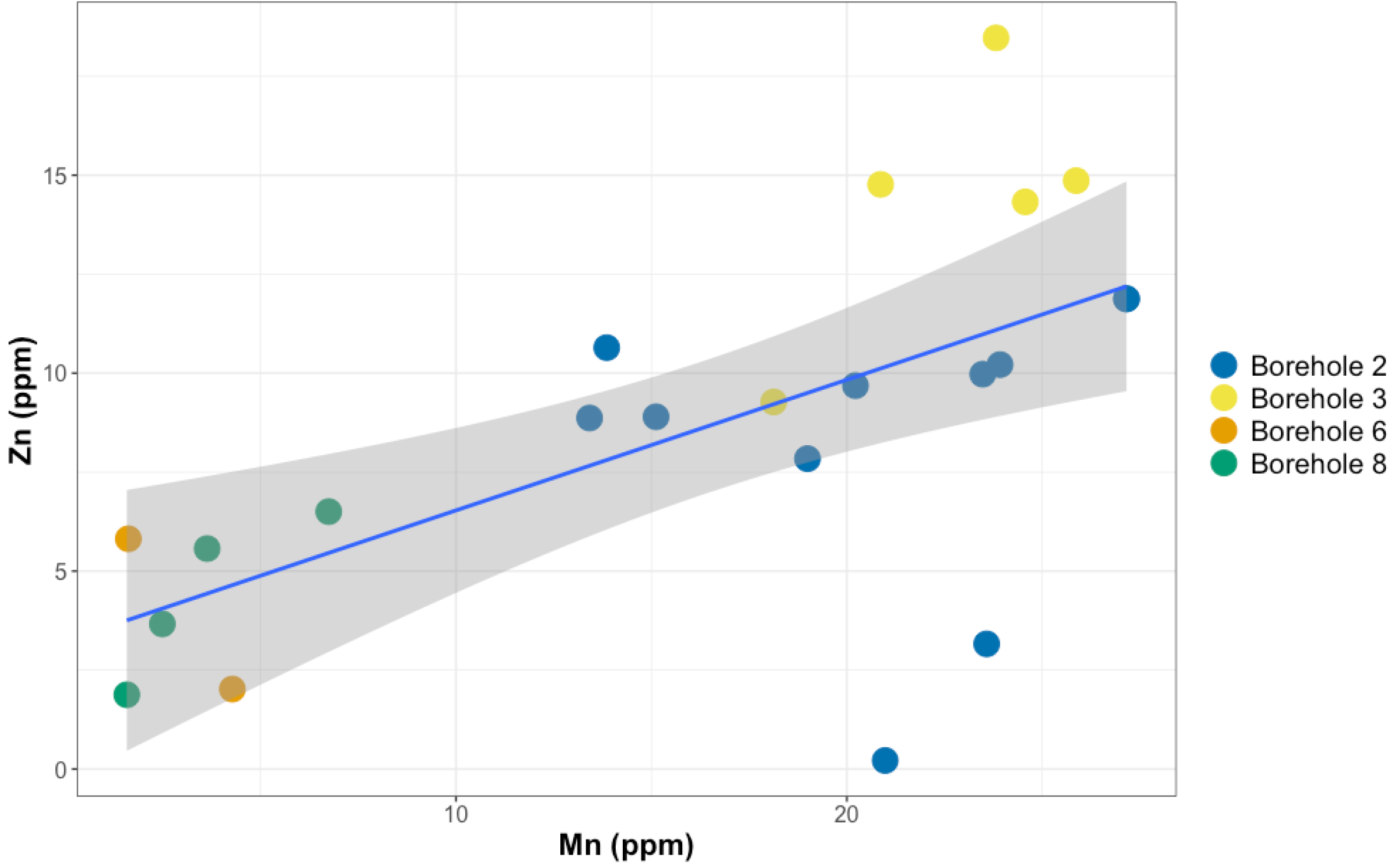
The calculated linear regression of ICP-AES data for Mn and Zn concentration within the borehole fluids. The linear regression line is plotted in blue with 95% confidence intervals shaded in grey (Adjusted R^2^ = 0.3643, F-statistic = 13.04, p = 0.00175). Samples are color coded from the borehole in which they were collected from.

**Table 1.**
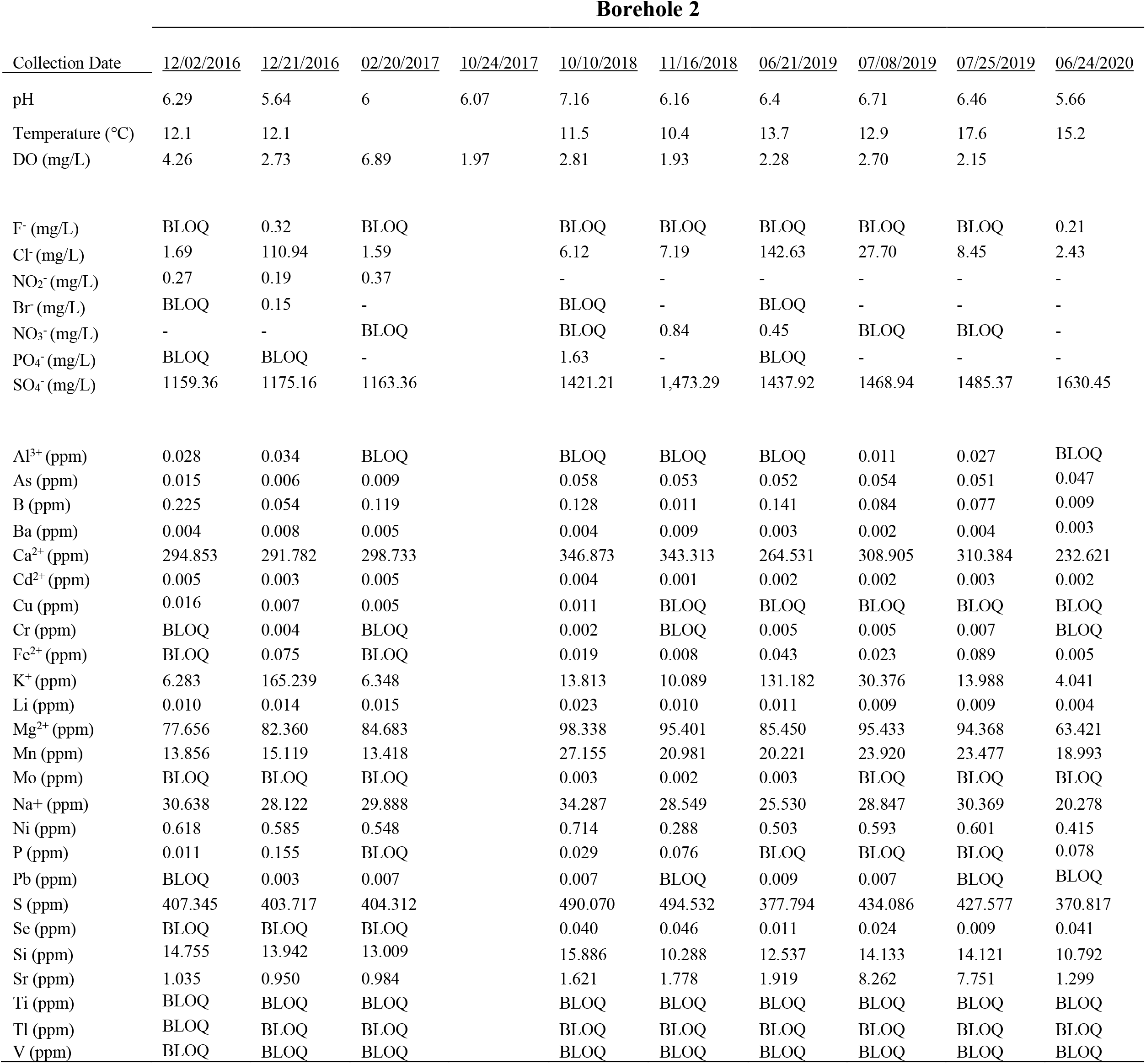
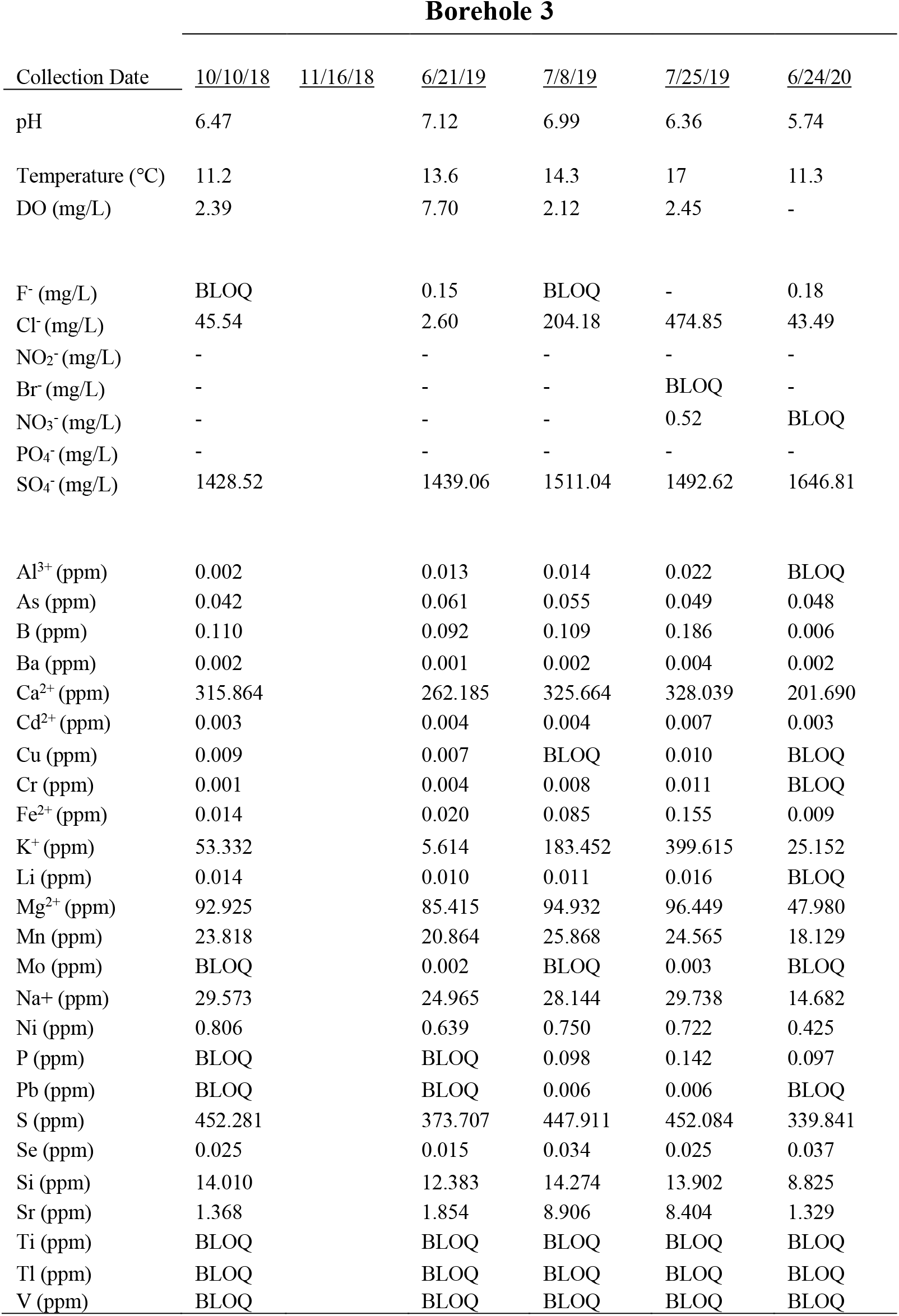
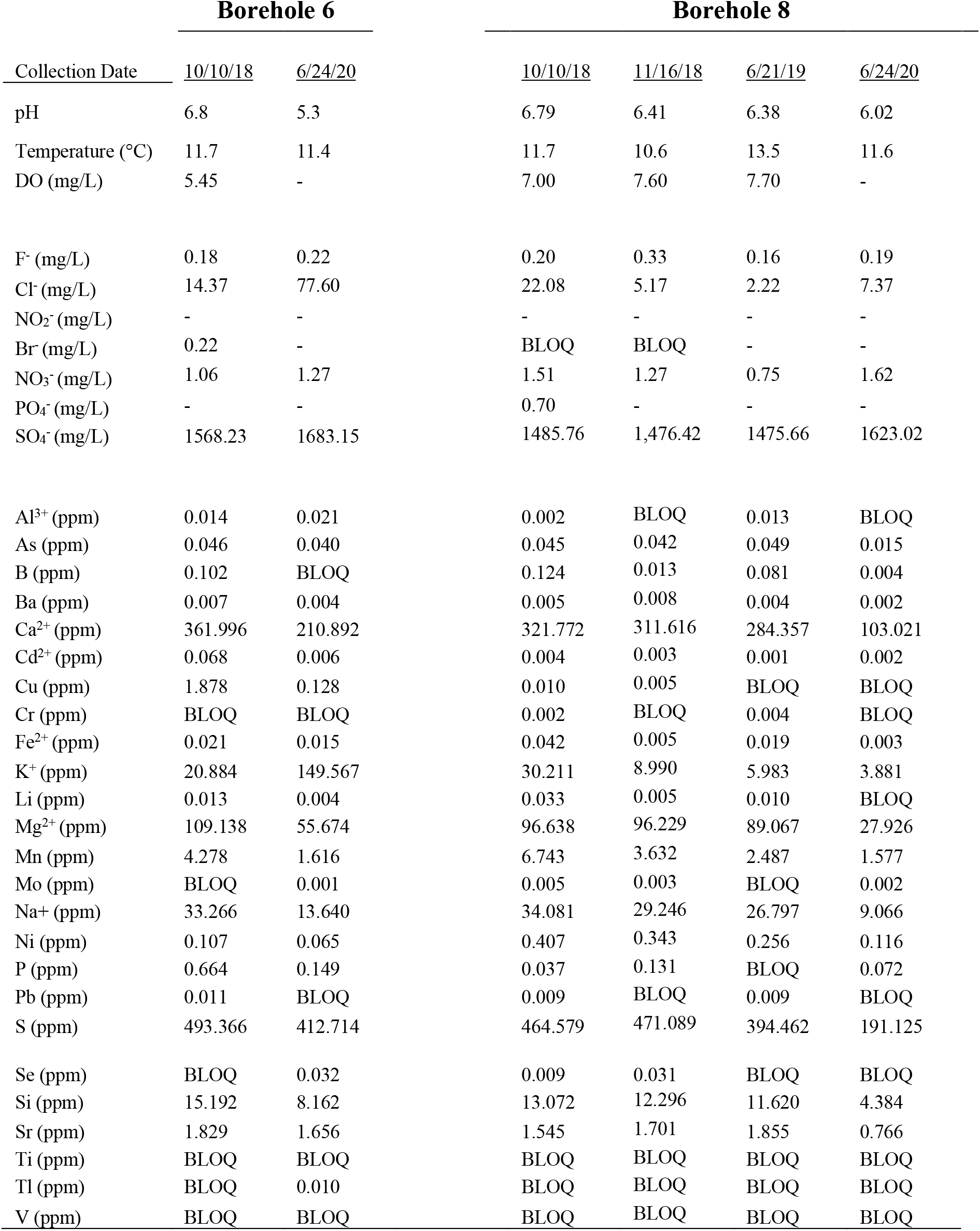
Measurements of major anions and cations in borehole fluids, as well as pH, temperature, and dissolved oxygen (DO). BLOQ = below limit of quantitiation, dashed lines indicate measurements were not taken.

Borehole 2
| Collection Date | <u>12/02/2016</u> | <u>12/21/2016</u> | <u>02/20/2017</u> | <u>10/24/2017</u> | <u>10/10/2018</u> | <u>11/16/2018</u> | <u>06/21/2019</u> | <u>07/08/2019</u> | <u>07/25/2019</u> | <u>06/24/2020</u> |
| --- | --- | --- | --- | --- | --- | --- | --- | --- | --- | --- |
| pH | 6.29 | 5.64 | 6 | 6.07 | 7.16 | 6.16 | 6.4 | 6.71 | 6.46 | 5.66 |
| Temperature (°C) | 12.1 | 12.1 |  |  | 11.5 | 10.4 | 13.7 | 12.9 | 17.6 | 15.2 |
| DO (mg/L) | 4.26 | 2.73 | 6.89 | 1.97 | 2.81 | 1.93 | 2.28 | 2.70 | 2.15 |  |
| F<sup>-</sup> (mg/L) | BLOQ | 0.32 | BLOQ |  | BLOQ | BLOQ | BLOQ | BLOQ | BLOQ | 0.21 |
| Cl<sup>-</sup> (mg/L) | 1.69 | 110.94 | 1.59 |  | 6.12 | 7.19 | 142.63 | 27.70 | 8.45 | 2.43 |
| NO<sub>2</sub><sup>-</sup> (mg/L) | 0.27 | 0.19 | 0.37 |  | - | - | - | - | - | - |
| Br<sup>-</sup> (mg/L) | BLOQ | 0.15 | - |  | BLOQ | - | BLOQ | - | - | - |
| NO<sub>3</sub><sup>-</sup> (mg/L) | - | - | BLOQ |  | BLOQ | 0.84 | 0.45 | BLOQ | BLOQ | - |
| PO<sub>4</sub><sup>-</sup> (mg/L) | BLOQ | BLOQ | - |  | 1.63 | - | BLOQ | - | - | - |
| SO<sub>4</sub><sup>-</sup> (mg/L) | 1159.36 | 1175.16 | 1163.36 |  | 1421.21 | 1,473.29 | 1437.92 | 1468.94 | 1485.37 | 1630.45 |
| Al<sup>3+</sup> (ppm) | 0.028 | 0.034 | BLOQ |  | BLOQ | BLOQ | BLOQ | 0.011 | 0.027 | BLOQ |
| As (ppm) | 0.015 | 0.006 | 0.009 |  | 0.058 | 0.053 | 0.052 | 0.054 | 0.051 | 0.047 |
| B (ppm) | 0.225 | 0.054 | 0.119 |  | 0.128 | 0.011 | 0.141 | 0.084 | 0.077 | 0.009 |
| Ba (ppm) | 0.004 | 0.008 | 0.005 |  | 0.004 | 0.009 | 0.003 | 0.002 | 0.004 | 0.003 |
| Ca<sup>2+</sup> (ppm) | 294.853 | 291.782 | 298.733 |  | 346.873 | 343.313 | 264.531 | 308.905 | 310.384 | 232.621 |
| Cd<sup>2+</sup> (ppm) | 0.005 | 0.003 | 0.005 |  | 0.004 | 0.001 | 0.002 | 0.002 | 0.003 | 0.002 |
| Cu (ppm) | 0.016 | 0.007 | 0.005 |  | 0.011 | BLOQ | BLOQ | BLOQ | BLOQ | BLOQ |
| Cr (ppm) | BLOQ | 0.004 | BLOQ |  | 0.002 | BLOQ | 0.005 | 0.005 | 0.007 | BLOQ |
| Fe<sup>2+</sup> (ppm) | BLOQ | 0.075 | BLOQ |  | 0.019 | 0.008 | 0.043 | 0.023 | 0.089 | 0.005 |
| K<sup>+</sup> (ppm) | 6.283 | 165.239 | 6.348 |  | 13.813 | 10.089 | 131.182 | 30.376 | 13.988 | 4.041 |
| Li (ppm) | 0.010 | 0.014 | 0.015 |  | 0.023 | 0.010 | 0.011 | 0.009 | 0.009 | 0.004 |
| Mg<sup>2+</sup> (ppm) | 77.656 | 82.360 | 84.683 |  | 98.338 | 95.401 | 85.450 | 95.433 | 94.368 | 63.421 |
| Mn (ppm) | 13.856 | 15.119 | 13.418 |  | 27.155 | 20.981 | 20.221 | 23.920 | 23.477 | 18.993 |
| Mo (ppm) | BLOQ | BLOQ | BLOQ |  | 0.003 | 0.002 | 0.003 | BLOQ | BLOQ | BLOQ |
| Na<sup>+</sup> (ppm) | 30.638 | 28.122 | 29.888 |  | 34.287 | 28.549 | 25.530 | 28.847 | 30.369 | 20.278 |
| Ni (ppm) | 0.618 | 0.585 | 0.548 |  | 0.714 | 0.288 | 0.503 | 0.593 | 0.601 | 0.415 |
| P (ppm) | 0.011 | 0.155 | BLOQ |  | 0.029 | 0.076 | BLOQ | BLOQ | BLOQ | 0.078 |
| Pb (ppm) | BLOQ | 0.003 | 0.007 |  | 0.007 | BLOQ | 0.009 | 0.007 | BLOQ | BLOQ |
| S (ppm) | 407.345 | 403.717 | 404.312 |  | 490.070 | 494.532 | 377.794 | 434.086 | 427.577 | 370.817 |
| Se (ppm) | BLOQ | BLOQ | BLOQ |  | 0.040 | 0.046 | 0.011 | 0.024 | 0.009 | 0.041 |
| Si (ppm) | 14.755 | 13.942 | 13.009 |  | 15.886 | 10.288 | 12.537 | 14.133 | 14.121 | 10.792 |
| Sr (ppm) | 1.035 | 0.950 | 0.984 |  | 1.621 | 1.778 | 1.919 | 8.262 | 7.751 | 1.299 |
| Ti (ppm) | BLOQ | BLOQ | BLOQ |  | BLOQ | BLOQ | BLOQ | BLOQ | BLOQ | BLOQ |
| Tl (ppm) | BLOQ | BLOQ | BLOQ |  | BLOQ | BLOQ | BLOQ | BLOQ | BLOQ | BLOQ |
| V (ppm) | BLOQ | BLOQ | BLOQ |  | BLOQ | BLOQ | BLOQ | BLOQ | BLOQ | BLOQ |

Borehole 3
| Collection Date | 10/10/18 | 11/16/18 | 6/21/19 | 7/8/19 | 7/25/19 | 6/24/20 |
| --- | --- | --- | --- | --- | --- | --- |
| pH | 6.47 |  | 7.12 | 6.99 | 6.36 | 5.74 |
| Temperature (°C) | 11.2 |  | 13.6 | 14.3 | 17 | 11.3 |
| DO (mg/L) | 2.39 |  | 7.70 | 2.12 | 2.45 | - |
| F<sup>-</sup> (mg/L) | BLOQ |  | 0.15 | BLOQ | - | 0.18 |
| Cl<sup>-</sup> (mg/L) | 45.54 |  | 2.60 | 204.18 | 474.85 | 43.49 |
| NO<sub>2</sub><sup>-</sup> (mg/L) | - |  | - | - | - | - |
| Br<sup>-</sup> (mg/L) | - |  | - | - | BLOQ | - |
| NO<sub>3</sub><sup>-</sup> (mg/L) | - |  | - | - | 0.52 | BLOQ |
| PO<sub>4</sub><sup>-</sup> (mg/L) | - |  | - | - | - | - |
| SO<sub>4</sub><sup>-</sup> (mg/L) | 1428.52 |  | 1439.06 | 1511.04 | 1492.62 | 1646.81 |
| Al<sup>3+</sup> (ppm) | 0.002 |  | 0.013 | 0.014 | 0.022 | BLOQ |
| As (ppm) | 0.042 |  | 0.061 | 0.055 | 0.049 | 0.048 |
| B (ppm) | 0.110 |  | 0.092 | 0.109 | 0.186 | 0.006 |
| Ba (ppm) | 0.002 |  | 0.001 | 0.002 | 0.004 | 0.002 |
| Ca<sup>2+</sup> (ppm) | 315.864 |  | 262.185 | 325.664 | 328.039 | 201.690 |
| Cd<sup>2+</sup> (ppm) | 0.003 |  | 0.004 | 0.004 | 0.007 | 0.003 |
| Cu (ppm) | 0.009 |  | 0.007 | BLOQ | 0.010 | BLOQ |
| Cr (ppm) | 0.001 |  | 0.004 | 0.008 | 0.011 | BLOQ |
| Fe<sup>2+</sup> (ppm) | 0.014 |  | 0.020 | 0.085 | 0.155 | 0.009 |
| K<sup>+</sup> (ppm) | 53.332 |  | 5.614 | 183.452 | 399.615 | 25.152 |
| Li (ppm) | 0.014 |  | 0.010 | 0.011 | 0.016 | BLOQ |
| Mg<sup>2+</sup> (ppm) | 92.925 |  | 85.415 | 94.932 | 96.449 | 47.980 |
| Mn (ppm) | 23.818 |  | 20.864 | 25.868 | 24.565 | 18.129 |
| Mo (ppm) | BLOQ |  | 0.002 | BLOQ | 0.003 | BLOQ |
| Na<sup>+</sup> (ppm) | 29.573 |  | 24.965 | 28.144 | 29.738 | 14.682 |
| Ni (ppm) | 0.806 |  | 0.639 | 0.750 | 0.722 | 0.425 |
| P (ppm) | BLOQ |  | BLOQ | 0.098 | 0.142 | 0.097 |
| Pb (ppm) | BLOQ |  | BLOQ | 0.006 | 0.006 | BLOQ |
| S (ppm) | 452.281 |  | 373.707 | 447.911 | 452.084 | 339.841 |
| Se (ppm) | 0.025 |  | 0.015 | 0.034 | 0.025 | 0.037 |
| Si (ppm) | 14.010 |  | 12.383 | 14.274 | 13.902 | 8.825 |
| Sr (ppm) | 1.368 |  | 1.854 | 8.906 | 8.404 | 1.329 |
| Ti (ppm) | BLOQ |  | BLOQ | BLOQ | BLOQ | BLOQ |
| Tl (ppm) | BLOQ |  | BLOQ | BLOQ | BLOQ | BLOQ |
| V (ppm) | BLOQ |  | BLOQ | BLOQ | BLOQ | BLOQ |

**Table 1.** Measurements of major anions and cations in borehole fluids, as well as pH, temperature, and dissolved oxygen (DO). BLOQ = below limit of quantitation, dashed lines indicate measurements were not taken.
|  | **Borehole 6** |  | **Borehole 8** |  |  |  |
| --- | --- | --- | --- | --- | --- | --- |
| <u>Collection Date</u> | <u>10/10/18</u> | <u>6/24/20</u> | <u>10/10/18</u> | <u>11/16/18</u> | <u>6/21/19</u> | <u>6/24/20</u> |
| pH | 6.8 | 5.3 | 6.79 | 6.41 | 6.38 | 6.02 |
| Temperature (°C) | 11.7 | 11.4 | 11.7 | 10.6 | 13.5 | 11.6 |
| DO (mg/L) | 5.45 | - | 7.00 | 7.60 | 7.70 | - |
| F<sup>-</sup> (mg/L) | 0.18 | 0.22 | 0.20 | 0.33 | 0.16 | 0.19 |
| Cl<sup>-</sup> (mg/L) | 14.37 | 77.60 | 22.08 | 5.17 | 2.22 | 7.37 |
| NO<sub>2</sub><sup>-</sup> (mg/L) | - | - | - | - | - | - |
| Br<sup>-</sup> (mg/L) | 0.22 | - | BLOQ | BLOQ | - | - |
| NO<sub>3</sub><sup>-</sup> (mg/L) | 1.06 | 1.27 | 1.51 | 1.27 | 0.75 | 1.62 |
| PO<sub>4</sub><sup>-</sup> (mg/L) | - | - | 0.70 | - | - | - |
| SO<sub>4</sub><sup>-</sup> (mg/L) | 1568.23 | 1683.15 | 1485.76 | 1,476.42 | 1475.66 | 1623.02 |
| Al<sup>3+</sup> (ppm) | 0.014 | 0.021 | 0.002 | BLOQ | 0.013 | BLOQ |
| As (ppm) | 0.046 | 0.040 | 0.045 | 0.042 | 0.049 | 0.015 |
| B (ppm) | 0.102 | BLOQ | 0.124 | 0.013 | 0.081 | 0.004 |
| Ba (ppm) | 0.007 | 0.004 | 0.005 | 0.008 | 0.004 | 0.002 |
| Ca<sup>2+</sup> (ppm) | 361.996 | 210.892 | 321.772 | 311.616 | 284.357 | 103.021 |
| Cd<sup>2+</sup> (ppm) | 0.068 | 0.006 | 0.004 | 0.003 | 0.001 | 0.002 |
| Cu (ppm) | 1.878 | 0.128 | 0.010 | 0.005 | BLOQ | BLOQ |
| Cr (ppm) | BLOQ | BLOQ | 0.002 | BLOQ | 0.004 | BLOQ |
| Fe<sup>2+</sup> (ppm) | 0.021 | 0.015 | 0.042 | 0.005 | 0.019 | 0.003 |
| K<sup>+</sup> (ppm) | 20.884 | 149.567 | 30.211 | 8.990 | 5.983 | 3.881 |
| Li (ppm) | 0.013 | 0.004 | 0.033 | 0.005 | 0.010 | BLOQ |
| Mg<sup>2+</sup> (ppm) | 109.138 | 55.674 | 96.638 | 96.229 | 89.067 | 27.926 |
| Mn (ppm) | 4.278 | 1.616 | 6.743 | 3.632 | 2.487 | 1.577 |
| Mo (ppm) | BLOQ | 0.001 | 0.005 | 0.003 | BLOQ | 0.002 |
| Na<sup>+</sup> (ppm) | 33.266 | 13.640 | 34.081 | 29.246 | 26.797 | 9.066 |
| Ni (ppm) | 0.107 | 0.065 | 0.407 | 0.343 | 0.256 | 0.116 |
| P (ppm) | 0.664 | 0.149 | 0.037 | 0.131 | BLOQ | 0.072 |
| Pb (ppm) | 0.011 | BLOQ | 0.009 | BLOQ | 0.009 | BLOQ |
| S (ppm) | 493.366 | 412.714 | 464.579 | 471.089 | 394.462 | 191.125 |
| Se (ppm) | BLOQ | 0.032 | 0.009 | 0.031 | BLOQ | BLOQ |
| Si (ppm) | 15.192 | 8.162 | 13.072 | 12.296 | 11.620 | 4.384 |
| Sr (ppm) | 1.829 | 1.656 | 1.545 | 1.701 | 1.855 | 0.766 |
| Ti (ppm) | BLOQ | BLOQ | BLOQ | BLOQ | BLOQ | BLOQ |
| Tl (ppm) | BLOQ | 0.010 | BLOQ | BLOQ | BLOQ | BLOQ |
| V (ppm) | BLOQ | BLOQ | BLOQ | BLOQ | BLOQ | BLOQ |

**Figure 3** illustrates ∂^18^O and ∂^2^H values for borehole fluids compared to the Global Meteoric Water Line (GMWL). Samples are plotted above the GMWL with slightly enriched ∂^2^H values. Edgar Mine is located at greater elevation and drier climate, which often causes enriched ∂^2^H values of meteoric water as a result from the rainout effect [21]. Meteoric water appears the most logical source of fluid for the boreholes also given the rate of fluid recharge and elevation of the mine, disregarding the notion for alternative groundwater sources feeding the boreholes.

**Figure 3.**
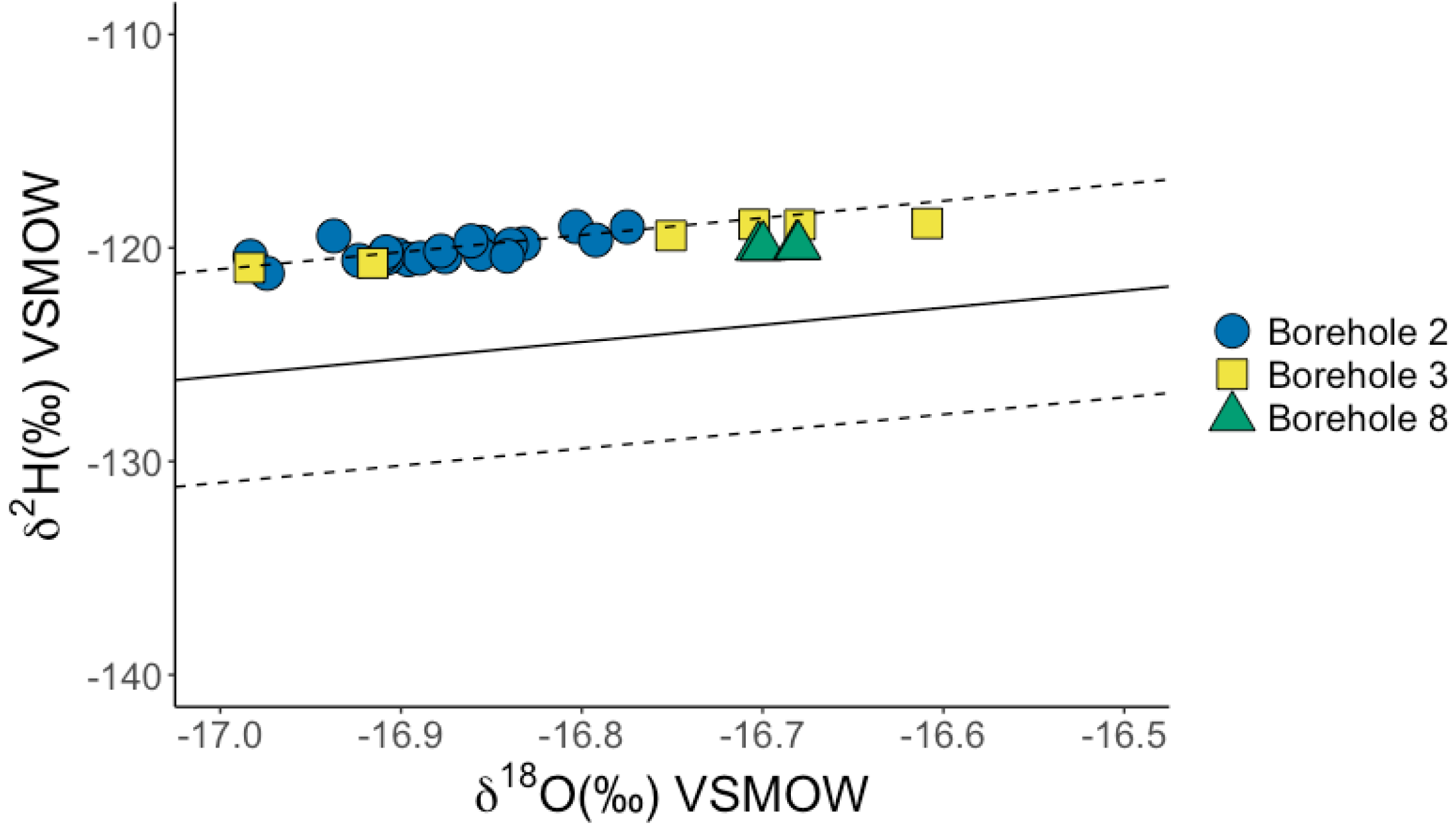
The ∂^18^O and ∂^2^H signatures for borehole fluids collected at Edgar Mine. The solid line indicates the trendline for the global meteoric water line (GMWL) (∂^2^H = 8 x ∂^18^O + 10) while the dashed lines represent ± 5‰ trendline amendments. There appears to be no flux in isotopic values from each borehole, indicating borehole fluids are likely meteoric water slightly enriched in ∂^2^H, likely due to the rainout effect as a result of the location of Edgar Mine. This data demonstrates there are no adverse processes altering the fluid isotope chemistry, nor is there an alternate source for the fluids percolating throughout the boreholes.

### Microbial Community Analysis

16S rRNA gene sequence analyses were performed on filtered water to determine the microbial community composition within each borehole. Several samples from each borehole were taken to contrast composition over time. The fluids were dominated by *Bacteria* (97.2%) with only a small presence of *Archaea* (2.8%) belonging predominantly to Borehole 2. All boreholes were collectively dominated by the phylum *Proteobacteria* (39.2%) and also contained sequences for divisions of *Firmicutes* (15.7%), indicating anaerobic conditions, and *Bacteroidota* (13.3%) known to represent a diverse group microorganisms.

Borehole 2 contained the presence of microbiota commonly associated in subsurface hosted systems representative of mining or metal-rich environments. The presence of *Gallionellaceae* reflects the greater concentrations of Mn and Fe from the metal-rich fluids [22, 23]. Sequence analysis identified *Acidiferrobacteraceae* and *Desulfobacteriaceae*, which contain known species of sulfur oxidizers and reducers, respectively [24, 25]. The large abundance of SO_4_-within Borehole 2 fluid suggests sulfur metabolisms are a dominant mechanism of microorganisms inhabiting the rock-equilibrated fluid. There is a stark contrast in microbial composition demonstrated from the last two sampling periods, after 948 days have passed into the experiment (**Figure 4**). The abundance of *Proteobacteria* drops sharply while the abundance of *Firmicutes* increases. While almost undetected in the majority of timepoints collected from Borehole 2, *Nanoarchaeota* sharply increase in abundance alongside *Firmicutes* making up 15.7% of the total microbial composition of the borehole fluid from the most recently collected sample. Similar fluctuations in microbial composition are demonstrated in Borehole 3 where the abundance of *Firmicutes* rises and falls inversely with *Proteobacteria* and *Bacteriodetes* abundance (**Figure 5**). This is represented by the inclusion and abundance of *Desulfosprosinus,* a strictly anaerobic sulfate-reducing bacteria and member of the *Firmicutes* phyla, observed in Borehole 2 [25]. When Borehole 3 is dominated by *Firmicutes*, the main genus level endmember identifies as *Sulfurifustis* indicating a role for sulfur-oxidizing metabolisms [24]. The drop in *Firmicutes* abundance results in greater diversification of microbial community composition within the borehole. The borehole becomes saturated with known aerobic microorganisms capable of a diverse array of metabolisms. It appears that more abundant members are capable of adapting to the more metal-rich fluids similar to Borehole 2. *SR-FBR-L83* is observed to respire electron acceptors such as Fe(III) and NO_2_-, while *Candidatus Nitroga* is capable of adapting to nitrite limited conditions and perform sulfur oxidizing metabolic functions [26]. Similarly, *Chelatococcus* requires metal-chelating compounds to use as sole energy sources [27]. DNA sequences from Boreholes 2 and 3 reveal microbial community compositions supporting the presence of members capable of surviving in metal-rich fluids that have equilibrated within the boreholes.

**Figure 4.**
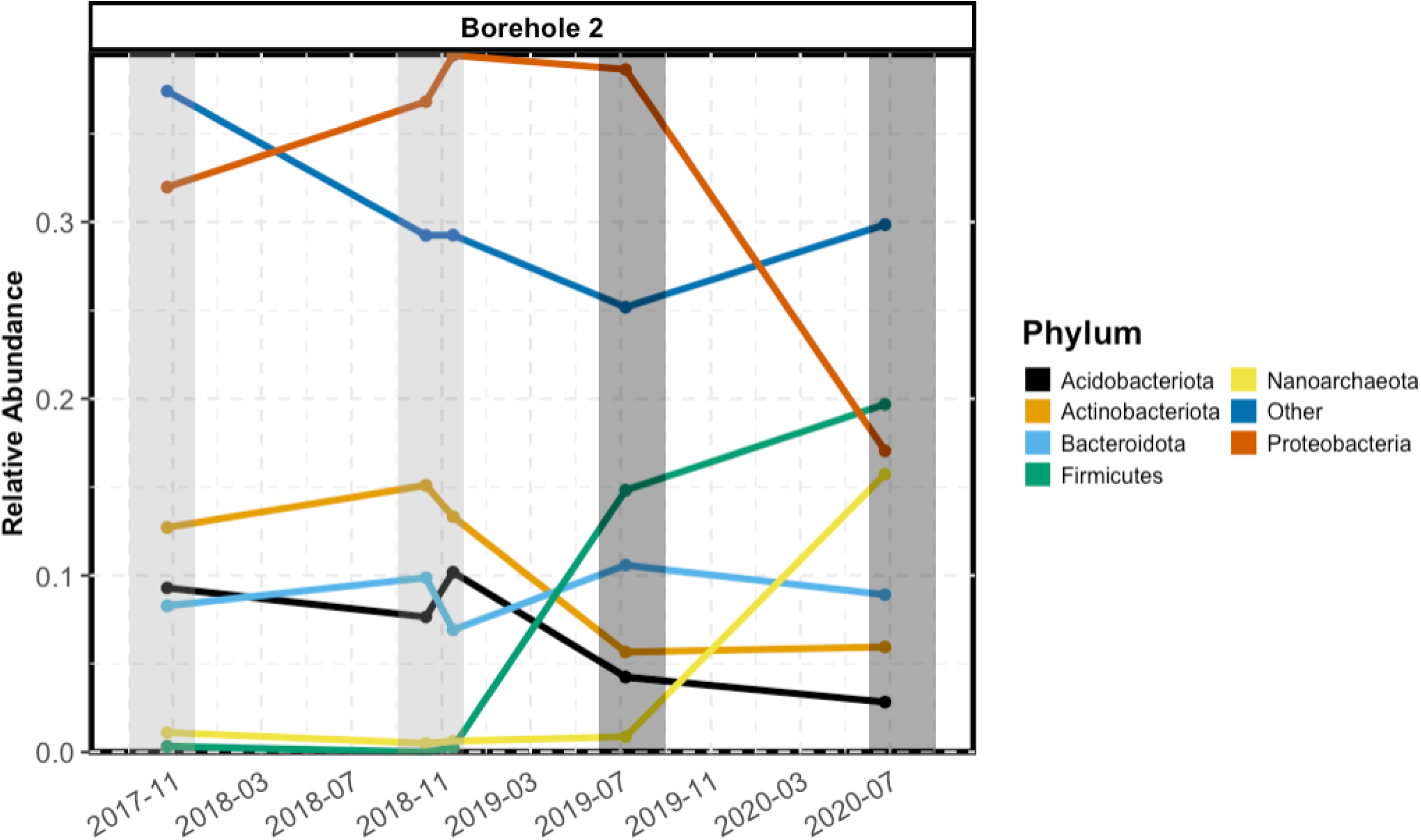
Relative abundances of major phylum level represented microorganisms collected from Borehole 2. Samples are represented as time points collected throughout the study, with the dates occurring within the span of the 4-year sampling period. The shaded areas indicate samples collected in the Fall (light grey), and samples collected in the Summer (dark grey) from meteorological seasonal classification. Seasons are scaled accordingly to demonstrate the time frame within a season samples were collected. Only the seasons in which samples were collected are highlighted. Phylum level sequences identified below 5% throughout the course of the experiment are summed together in the “Other” classification to reveal the relative abundance of the low abundance members of the microbial community within Borehole 2.

**Figure 5.**
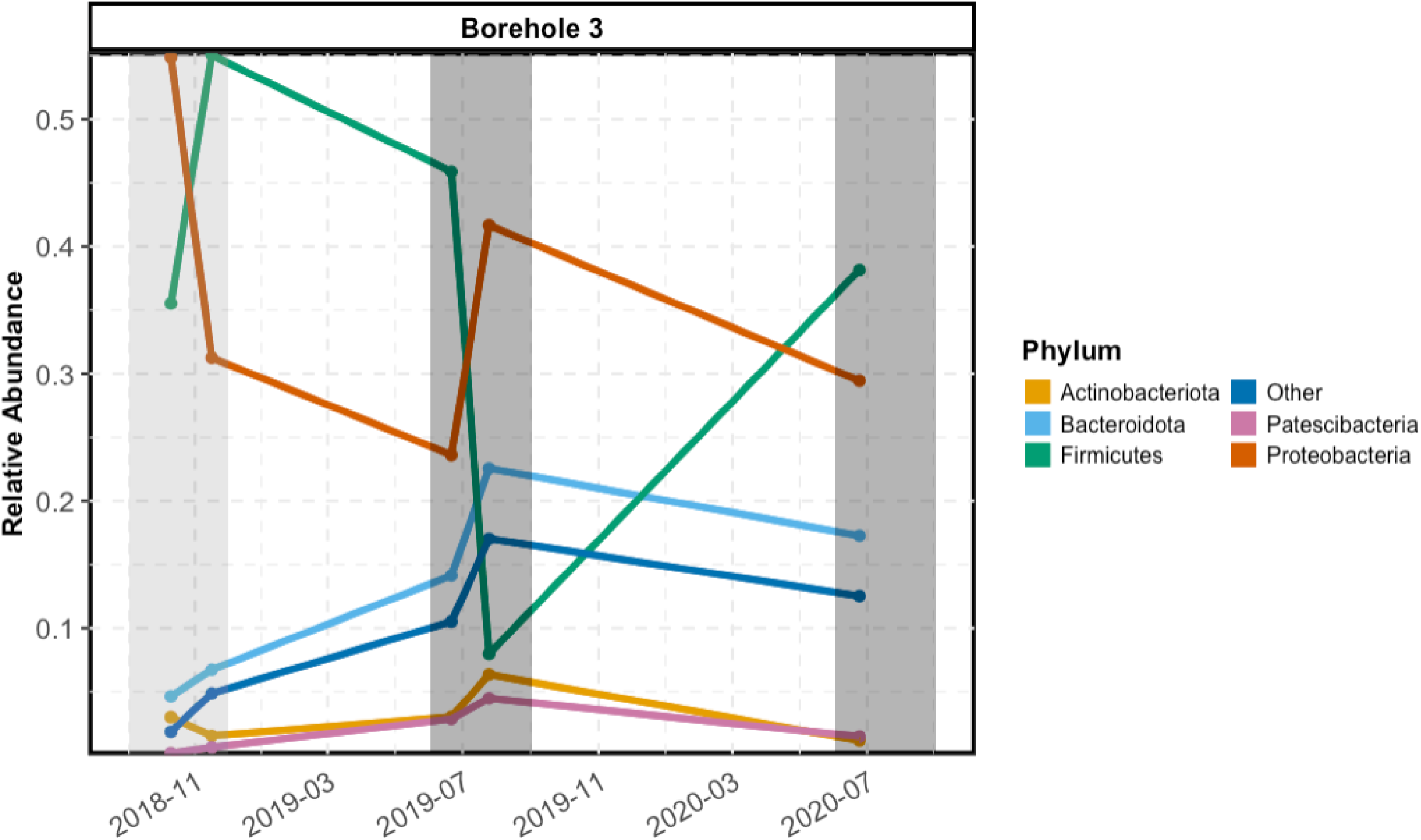
Relative abundances of major phylum level represented microorganisms collected from Borehole 3. Samples are represented over a shorter time scale of the sampling period of the experiment due to the lack of consistent fluid recharge within the borehole. Similarly to Figure 4, the shaded areas indicate Fall (light grey) and Summer (dark grey), and scaled accordingly within the time frame of the study when samples were collected. The “Other” classification makes up the sequences identified below 5% throughout the course of the experiment within Borehole 3.

Comparatively, Boreholes 6 and 8 do not display a temporal influence in microbial community composition and abundances. The lack of temporal evaluation of these two boreholes is owed to the unpredictability of fluid recharge. Despite the opportunity to retrieve additional samples, Boreholes 6 and 8 maintain microbial compositions suggesting more soil influenced communities (**Figure 6)**. This is highlighted by fluids more deplete in metal concentrations, yet these boreholes display the only detectable concentrations of nitrate. Compared to Boreholes 2 and 3, Boreholes 6 and 8 have a greater concentration of DO providing a more aerobic environment. Borehole 6 DNA sequencing reveals genus level microbial members commonly associated with soil environments including *Afipia*, *Sediminibacterium*, and *Sphingobium* [28–30]. The inclusion of soil-derived microorganisms could indicate a more direct link to the surface by way of an undefined fracture system. Similarly, Borehole 8 encompasses a community defined by known soil originating microorganisms. *Aminobacter, Bradyrhizobium,* and *Candidatus Nitrosotenuis* are among known soil inhabiting genera possessing nitrogen fixing and reducing metabolic capabilities [31, 32]. The unexpected inclusion of *Polaromonas* in Borehole 8 may be a result of snowmelt acting as the meteoric source for fluid recharge [33, 34], which is congruent with isotopic findings that support meteoric water as the primary hydrologic source (Figure 3). Boreholes 6 and 8 demonstrate microbial community compositions more representative of soil environments, which indicate conditions favorable only to relict microorganisms capable of utilizing adaptable metabolisms with suppressed nutrient availability in the subsurface (**Figure 6**).

**Figure 6.**
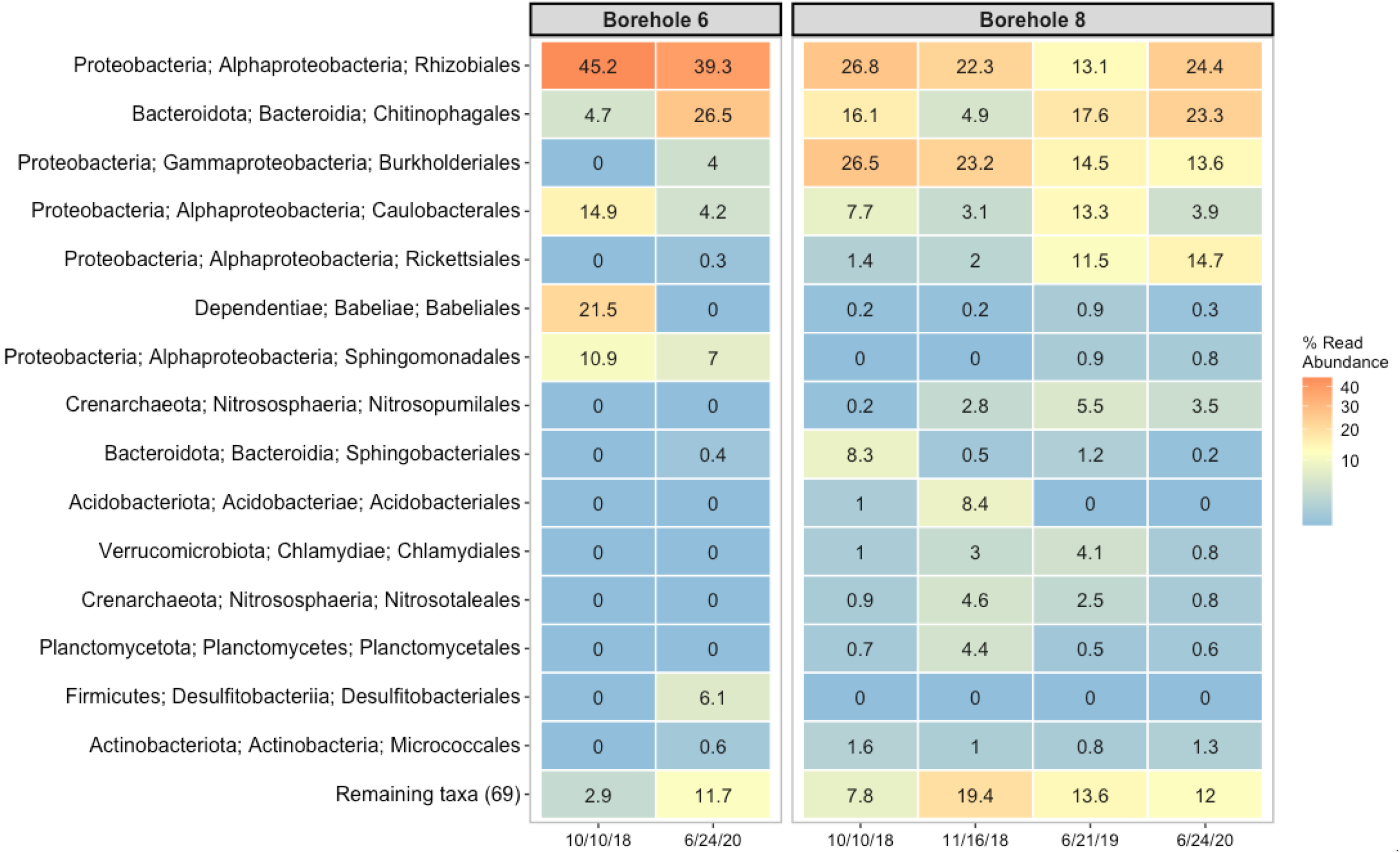
The top 15 order-level microorganisms within boreholes 6 and 8 as a function time over the course over the experiment. Each column indicates the sampling date in which a sample for the respective borehole was collected. Numeric values within the box indicate the percent read abundance, also represented as a heatmap through boxes colored by the numerical ranges represented from high (orange) to low (blue). The full phylogenetic lineage from phylum to class to order is depicted to give full coverage of the organism in abundance within each borehole.

Ordination via PCoA with a weighted Unifrac distance matrix was performed to determine statistical similarity of microbial communities from each borehole (**Figure 7**) [35]. Ordination suggests strong clustering as a result of grouping by where the borehole samples were collected. Additional soil samples were collected from the surface of the mine to confirm the distinctive compositions of borehole microbiota. An Adonis test was run on the dissimilarity matrix to verify sampling source as distinct and significant (R^2^ = 0.44, *p* = 0.001). Weighted Unifrac distance matrices were also tested by Adonis for the number of days since the initial extraction of fluid from the boreholes, the number of days between sampling events, and the season samples were collected. Results demonstrated the number of days since the initial extraction period as significant (R^2^ = 0.38, *p* = 0.001) as well as seasonality (R^2^ = 0.36, *p* = 0.001), and the number of days between sampling events as modestly significant (R^2^ = 0.073, *p* = 0.047). These values suggest each borehole is host to unique microbial community compositions. The inherent covariance in our study between days since initial extraction and seasonality suggests further work is needed to decipher these two signals. Nonetheless, a clear shift in community composition occurs over time, predominantly in Boreholes 2 and 3. However, the source of this change requires additional sampling to account for unsampled seasons and how microbial diversity may change with even more time allowed to pass within the boreholes.

**Figure 7.**
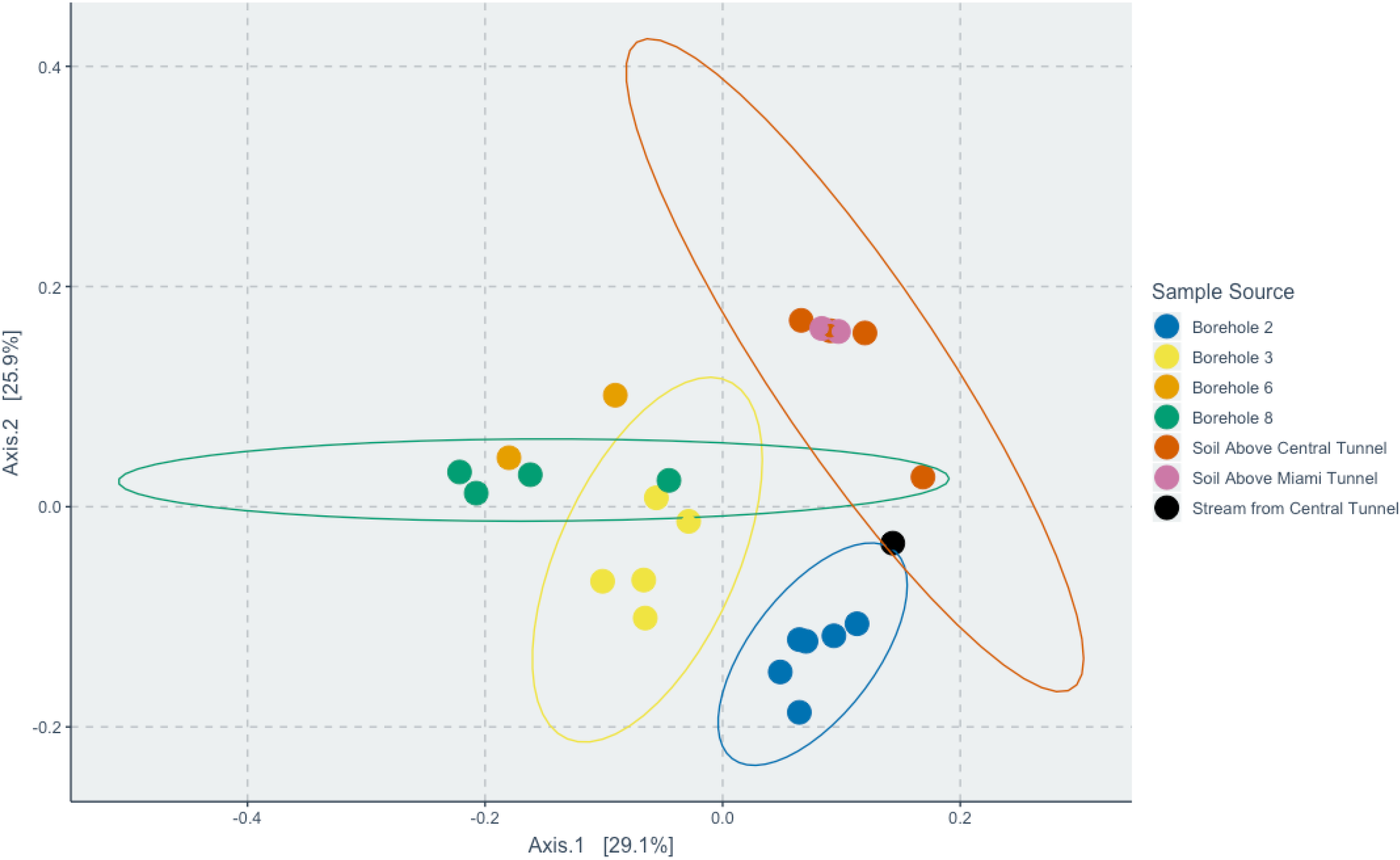
Principal coordinate analysis (PCoA) of weighted Unifrac distances between samples collected from borehole fluids and soil samples collected above Edgar Mine. Samples are color coded from the source they were collected from. Sample groupings with enough data were able to determine normal distribution ellipses. Differences between the samples collected from each borehole and the soil samples is significant as verified by an Adonis test (R^2^ = 0.45, p = < 0.001).

## Discussion

The near-surface subsurface biosphere links the roles of surface and hydrogeological interactions driving habitability for microorganisms in subsurface environments. We have presented this biological investigation at Edgar Mine, a near-surface experimental/demonstration mine owned by the Colorado School of Mines, to establish the conditions influencing microbiology within the shallow subsurface. This study presents a process for determining the microbiology of subsurface borehole fluids from an active mine, contributing to understanding surface and near-subsurface interaction dynamics. By including the temporal facet of sample collection, we were able to evaluate the evolution of borehole fluids overtime and how successive recharge events affect microbial community composition.

### Geochemical Dynamics

The chemistry of the subsurface fluids in the Edgar Mine reflects the hydrologic context as well as the extent of equilibration with the host rock. Due to the quick recharging of Borehole 2, it appears that residence time of meteoric water infiltrating the subsurface is quick (<14 days). No significant relationship between the amount of fluid collected and chemical composition of each borehole is apparent. Chemical differences between boreholes are minor, which suggests small scale variations in fluid flow paths defined by a convoluted fracture system within the host rock, at least in this near-subsurface environment. Concentrations of metals are likely the result of fluid-rock equilibration that may be unique to each borehole’s mineral inclusions. By extrapolation, this then has relevance whereby particular drill holes, mines sites, or access points are really detecting and capturing subsurface microbiology at precise screenshots within the subsurface with greater variability being present than is actually determined by these single slices of data. The fracture system in the Edgar Mine remains to be fully investigated, but requires more effort to decipher the link of surface influences into the subsurface and interconnectedness between the boreholes. In other subsurface geobiological datasets (both historical and future), caution should be taken in the application of ‘pin-prick’ biopsy observations to the whole of the below-ground.

∂^18^O and ∂^2^H values plotted along the GMWL support meteoric water as the fluid source for the boreholes. Without a source of water from the deep subsurface or ancient aquifer, Edgar Mine poses as a unique boundary at the surface and near-subsurface interface. Meteoric water transported from the surface allows initial investigation into how the subsurface interacts with surface derived fluids and how said fluid and boreholes evolve over time. The slightly enriched ∂^18^O and ∂^2^H values hold true to Edgar Mine’s inland location at elevation and drier climate. Minor shifts in ∂^18^O and ∂^2^H values reflect seasonality, but do not suggest any alteration of the fluid through supplementary subsurface processes [36].

The chemistry of Edgar Mine waters was further analyzed to understand biologically relevant chemical concentrations of borehole fluids and their effect on the composition of borehole microbial communities. Geochemistry of the borehole fluids varied weakly over the course of the experiment, which contrasts with heterogeneity in the composition of sequenced microbial communities. Instead, borehole geochemistry remained relatively consistent displaying neutral conditions with low concentrations of analytes for potential microbial metabolic functions. Additionally, the incubation of fluid between extraction periods does not seem to alter fluid geochemistry. Solute transport may not play as significant a role between the surface and near-subsurface influencing borehole fluids given the lack of geochemistry that would reflect soil origination, though additional analyses of dissolved organic carbon and nitrogen would aid in this inquiry. Nonetheless, in the more nutrient-limited deep subsurface, the movement of dilute geochemical analytes from above to below-ground is likely critical to recluse microbial metabolisms.

Inorganic nitrogen levels were low or depleted suggesting the near-subsurface boreholes of the Edgar Mine could be a nitrogen-limited system. Similarly, phosphate concentrations were below detection level and general phosphorus levels were extremely low, further suggesting a phosphorus-limited environment. One key factor driving potential microbial metabolisms and influencing fluid geochemistry is the elevated sulfate concentrations. Sulfate is highly concetrated in the borehole fluids and is likely a major component in sulfur reducing metabolisms [37, 38]. The positive correlation between Mn and Zn (**Figure 2**) demonstrates a relationship between the metals concentrations and the borehole fluid origin; which can be indicative of the host environment for microbial communities [39–41]. Boreholes 2 and 3 contain the greatest amounts of the metals, while Boreholes 6 and 8 contain the lowest concentrations. This gradient distinguishes the metal-rich fluids versus the more dilute fluids that can host diverging metabolic potentials.

### Borehole Microbial Community Analysis

Groundwater collected from packer installment allowed for fluid-rock equilibration spanning weeks to months between subsequent sampling events. While the undefined fracture system produces an unpredictable volume of fluid, both the rock and mineral composition appears to be unique to individual boreholes at the centimeter scale. The chemical gradient established within the more metal-rich boreholes is associated with a shift in community composition, juxtaposed by more dilute and aerobic boreholes. All boreholes are filled with similarly sourced meteoric water infiltrating the subsurface, yet each borehole displays a unique community composition as determined from PCoA weighted Unifrac analysis (**Figure 7**), and microbiota relative abundance investigations (Figures 4-6). Phylogenetic relatedness determined by PCoA analysis of borehole microbial communities suggests niche microcosms of microorganisms can establish themselves in boreholes located only centimeters to meters apart. Subsurface microbiology may be more sensitive to fluid-rock equilibrated systems at a finer spatial resolution than previously thought or understood. The host rock composition allowing for the dissolution of more metals or anions into fluids may be a key driver for supporting microbial habitability, as well as developing an aerobic/anaerobic environment. There are no discrete geochemical parameters that define microbial diversity observed in the boreholes, yet each borehole must provide unique electron donors and acceptors to support life. While dissolved inorganic carbon (DIC) and dissolved organic carbon (DOC) compounds were not investigated, carbon sources within each borehole likely influence microbial community composition. A model considering all carbon and electron donor and acceptor modulations that drive differences in our samples would be speculatory; nonetheless, subtle fluid variations in DOC and dissolved organic nitrogen (DON) likely assist in structuring the microbial heterogeneity that is unexplained by measured anions and cations, alone.

There appears to be an evolution of microbial community composition as the experiment progressed, reaching greater than 800 days. Previous studies have determined the effect of pre- and post-packer installment in establishing changes in microbial composition [42]. The installment of a packer into a borehole allows for fluid to equilibrate with the host rock and provide an opportunity for metals and other nutrients to become available to microbial communities. Similar studies have investigated boreholes of deep mine systems, yet not on such a frequent and lasting temporal scale or within a near-surface environment [17, 19]. The Edgar Mine boreholes are supplied by recurring recharge events from meteoric water, which may vary in geochemical composition when transported into the subsurface. This non-consistent recharge could play a role in determining the habitability of the near-surface subsurface biosphere.

Intriguingly, the time between sampling periods did not demonstrate a significant correlation to microbial community composition (**Figure S1**). This finding suggests defined time intervals between borehole fluid extractions does not dictate the composition of the microbial community within a borehole. Instead, the temporal influence is sourced from moderate changes over a greater period of time reflecting stages of evolution of the microbial community in borehole fluids. This finding highlights that microbial communities found in the shallow subsurface borehole fluids are not predictable based on the time between extractions. The season in which samples were collected also appears to display a strong correlation. This seasonal signal is difficult to discern and is biased from overlapping signal produced by the days since the initial fluid extraction. Nevertheless, seasonality may introduce an interesting avenue of further examination to determine the source of groundwater and how this may alter the near-subsurface microbial community [43, 44].

Snowfall may carry large amounts of biodiversity, especially through storm events, which could introduce nonindigenous microorganisms into the subsurface [45]. Transportation of microbial members through snowmelt has been speculated at the Henderson Mine near Empire, CO from the presence of *Chloroflexi* phylotypes [42]. The presence of *Polaromonas* in Boreholes 3 and 8 offers the same insight given the ≥98% identity match through BLAST search of sequences for a species isolated from an ice core [33]. The role of snowmelt transportation into the near subsurface requires further investigation to elucidate the role of meltwater and other seasonal hydrological infiltration into the subsurface as a mode for microbial transportation. Additionally, the role of water extraction on overlying litter/soil communities with subsequent seepage into the subsurface could further explain source and seasonal variation in near-surface subsurface environments. The role of hydrology intercepting the subsurface and introducing microbial diversity has direct relevance towards agriculture and home use that rely on near-surface water wells. Hydrological transport of microbial communities is also tied to the ever-present effects of climate change, which—through the mechanism of ground infiltration—will inevitably play a role in altering the geochemical and microbial composition of the subsurface, a concept likely also relevant to natural caves on Earth.

The presence of non-endemic microbial members is indicated by gradients in geochemistry across boreholes for aerobic conditions with more dilute metals concentrations. Boreholes 6 and 8 culminate an assembly of microorganisms typified by soil-originating sources (**Figure 6**). It should be noted that many of these organisms require aerobic conditions which could have been established from installment of the packer system [31]. When considering the microbial communities of these borehole fluids from the Edgar Mine, there is potential for contamination from operations within the mine. Specifically, the role of human activity, drilling fluids, and machinery used to drill the boreholes and install the packers in this demonstration and teaching mine cannot be ignored. This can make particular members of the microbial community difficult to determine if they are indigenous to the subsurface borehole ecosystems or a result of mining operations [46]. Nonetheless and despite this caveat, the composition of the microbial communities observed in Boreholes 6 and 8 may suggest a surface-influenced subsurface community. Of particular note, these two boreholes demonstrate the only quantifiable concentrations of nitrogen in the form of nitrate (**Table 1**). The presence of soil originating microbes with known nitrogen related metabolisms such as nitrification displayed by *Candidatus Nitrosenuis* may suggest any residual nitrogen compounds are quickly expended and are not left dissolved as inorganic fractions in the borehole fluid [32]. The presence of *Bradyrhizobium* further suggests the infiltration of soil derived microbes, as it is a known nitrogen fixing bacteria commonly found in plant soils [33]. Additional members like *Sphingobium* make it clear that the microbial communities observed in Boreholes 6 and 8 do not originate in the subsurface given the known metabolisms of phylotypes identified in the fluids [30]. Boreholes 6 and 8 contain microbial communities that suggest surface soil microorganisms were transported into the near-subsurface. This further indicates the interaction between the surface and subsurface, and how subsurface microbial communities can establish through surface infiltration. The development of surface-based communities in the subsurface remains to be fully explored, yet should press warning into determining true endemic members of the subsurface, in any subsurface location. The proximity of Boreholes 6 and 8 to Boreholes 2 and 3 illustrates how the spatial resolution for subsurface microbiology may be finer than we understand. The need for greater data collection emphasizes the potential of under-sampling over both spatial and temporal scales and reflects our lack of knowledge towards the true diversity and dynamic capabilities of microbial ecosystems in subsurface investigations.

The distinct differences in microbial community composition between boreholes is highlighted in the PCoA weighted Unifrac analysis. Boreholes 2 and 3 contain microbial members that suggest the presence of a more native community of microorganisms to the boreholes of the Edgar Mine. The presence of *Gallionellaceae* reflects the greater concentrations of metals in the fluids of Borehole 2 given the known Fe-oxidizing and potential Mn-oxidizing metabolic capabilities [23]. Similarly, the presence of genera *Sulfurifustis* and *Desulfosporosinus* indicate metabolic potentials for sulfur oxidation and sulfate reduction respectively [24, 25]. The prominence of sulfate would suggest that sulfur metabolisms play a strong role, however, lack of knowledge of dissolved organic or inorganic carbon compounds makes it difficult to decipher the metabolic pathways occurring in these boreholes [37, 38]. The unexpected increase in abundance of *Nanoarchaeota* remains inconclusive, yet intriguing regarding the role they play in the subsurface boreholes of the Edgar Mine. Given their known syntrophic relationships and connection to viral distribution, more work is needed to decipher who the host of this organism might be and what function is operating within this member of the subsurface [47]. Known syntrophs of *Nanoarchaeota* include members with sulfur metabolic capabilities in hyperthomophilic environments [48, 49]. The strong inclination for sulfur cycling in the Edgar Mine boreholes would suggest that the presence of *Nanoarchaeota* are not unlikely, however, requires more work to discover the mechanism behind their increase in abundance—including below-ground viral ecology not represented in this study. Inclusions of sulfide veins would offer direct microbe-mineral interactions to support sulfur oxidizing metabolisms [50]. It is difficult to discern the bioenergetics observed in Borehole 2, but given the low concentration of dissolved oxygen with available electron acceptors (eg., SO_4_- and Fe), it is feasible that anaerobic chemolithotrophy is a prominent metabolic process occurring in the metal rich boreholes [51]. Borehole 3 shares similar microbiota capable of using metal rich fluids or surfaces for their energy sources, or adapting metabolic abilities in the case of *Candidatus Nitroga* to limited nitrite conditions [52]. Presence of microbes respiring metals for electron donors and acceptors are incorporated in the fluid that has equilibrated with the host rock. The presence of these microbes in Boreholes 2 and 3 offers speculation for pre-existing biofilms that are equilibrating with the infiltrating meteoric water [53, 54]. Leaking water from unpacked boreholes has already produced visible discolorations along the walls of the mine (**Figure S2**). Brown and yellow discoloration with white precipitates suggest a prospective environment for biofilms with rich mineralization to develop. Further investigation into pre-existing biofilms could benefit from examining boreholes without leaking fluids. Installation of a packer and injecting sterilized fluid into the borehole could determine how the host rock equilibrates with the fluid and what microbiology may be native to the borehole [55, 56].

Tools for microbiology to aid in economic mining practices have yet to be fully investigated. Our study demonstrates how slight nuances in borehole microbiology can occur at centimeter to meter scales, and these measurable differences occur at a greater rate than differences in fluid geochemistry. Understanding how microbial community composition reflects mineralogy and potential ore formations could be a useful tool to aid the development of less invasive mining practices. High sulfate concentrations are reported within all of our boreholes, which suggests potential bio-leaching of low-grade ore and sulfide veins could be occurring and this would not be unexpected in Colorado [50,57,58]. Using molecular biological based tools to investigate and identify ores or resources of interest would pose a more environmentally friendly and economically sustainable method of resource detection [50, 59]. In conjunction, the ability to tie microbial diversity with predictive hydrological behaviors may be needed to demonstrate the connectivity of surface and subsurface interactions. Modeling parameters combined with molecular data could generate a powerful tool in understanding the path and composition of subsurface fluids.

The work we present here demonstrates the capability for the Edgar Mine to establish itself as a premier subsurface facility for microbiological experimentation. This is the first full molecular biological research experiment conducted at the Edgar Mine, an experimental-teaching-demonstration mine that was created to train people mining engineering practices, and this study unveils the complexity of boreholes within a spatial distance of centimeters to meters apart. Our work also begins to investigate the role of the shallow subsurface and the dynamic relationship it shares with surface influences into the near-subsurface. The ability to define how the subsurface biosphere interacts and is possibly shaped by surface infiltration still needs further investigation to contrast the deep biosphere. Understanding subsurface microbial diversity is an important tool for determining potential sources of mineralogy and ore deposits for less invasive mining practices, and will also be essential in unveiling biogeochemical cycling of the subsurface at locations all over the Earth and other planetary bodies.

## Methods

### Field Site

Fluids from boreholes previously drilled in the Edgar Experimental Mine were collected beginning in Fall of 2016 and extending through Summer of 2020. The boreholes were created as a function of mining and drilling demonstrations for teaching purposes by the Colorado School of Mines, who also own the facility and guide operations. Edgar Mine contains thousands of available boreholes, and more continue to be drilled as a part of training efforts for students at the Colorado School of Mines interested in mining practices. Edgar Mine is located in Idaho Springs, Colorado and acts as a repurposed gold, silver, lead, and copper mine for teaching and research initiatives incorporating mining engineering and safety. As a function of demonstration practices, boreholes were drilled into the walls of the site named “C-Right” (**Figure 1)**. Spatial resolution of the boreholes varies on the scale of centimeters to meters in distance of each other and differ greatly by orientation of drilling angle. The mine is composed of Precambrian rock, primarily by gneisses, where the specific field site of “C-Right” dominated by quartz-plagioclase gneiss and quartz-plagioclase-biotite gneiss [60].

It was observed that many of the boreholes were leaking fluid suggesting a complicated fracture system allowing for opportunistic boreholes to collect substantial amounts of fluid. This fluid serves the potential for many water-rock interactions fueling microbial habitability. Similarly, fluid buildup for extended periods of time can develop unique water-rock equilibration hosting niche microenvironments in separate boreholes for diverse microbial communities [58, 59]. In order to take advantage of the leaking fluid, boreholes were fitted with custom-built expandable packers at the mouth of the borehole to trap leaking fluid **(Figure S3)**. These fluids were then left to equilibrate with the rock and develop microbial communities on temporal scales of just over two weeks to over a year to assess temporal variability within sampled boreholes. The field site “C-Right” within the Mine is located approximately 500 feet below the surface. We propose meteoric water is the main source of fluid infiltration into the subsurface, following an undefined fracture system allowing certain boreholes to fill with greater amounts of fluid than others. Infiltration of surface fluids also possess potential to navigate towards the subsurface boreholes and influence both geochemical and microbial processes differentially over space and time.

### Sample Collection

Collection of fluid was accessed from the expandable packer at the mouth of each borehole to assess both geochemistry of the fluid and microbial community composition. A sterile syringe was used to draw fluid trapped behind the packer and passed through a 0.22 µm filter to collect biomass. Filters were immediately frozen on dry ice in the field and stored at − 20°C back in the lab until further extraction and processing. Upon initial extraction of borehole fluid, the preliminary 50 mL of fluid was used to record physical chemistry measurements of dissolved oxygen, pH, and temperature immediately using a daily-calibrated Hach multiparameter field meter (HQ40D, Hach, Inc., Loveland, Colorado). An additional 50 mL and up to 1 L of filtered fluid was collected in clean autoclave-sterilized glass vials for each borehole for downstream geochemical and water isotopic analyses. To contrast surface microbiology (a potential high-biomass origin of subsurface-infiltrating microbiota), soil samples were collected above the “C-Right” field site and put on dry-ice in the field and stored at −20°C until further processing. All further analysis and preparation were conducted in the laboratory. Samples were collected over various time scales in order to assess temporal variability of microbial community composition and geochemical fluctuation. Borehole fluid was allowed to incubate for as short as 17 days and as long as 351 days within certain boreholes. Due to the unpredictable nature of the fracture system, some boreholes collected greater quantities of infiltrated fluid compared to others and, thus, were subjected to more frequent sampling opportunities. Pre-determined incubation periods (2 weeks) were used to assess how microbial community composition may alter from human-chosen time intervals. Similarly, the amount of time passed since the initial fluid extraction was also recorded to observe overall temporal effects shifting microbial diversity.

### Chemical Analysis

Filtrate was collected and analyzed for major ion concentrations through ion chromatography (IC) and inductively-coupled plasma atomic emission spectroscopy (ICP-AES) spectroscopy. IC analyses were run on a Dionex ICS-900 IC system while ICP-AES was performed on a Perkin-Elmer 8300 ICP-AES. ICP samples were acidified with 3-5 drops of 10% nitric acid with a total volume of 12 mL for sample submission. Additional fluid collected from boreholes was run on a Picarro Water Isotope Analyzer (D/H and ^18^O/^16^O) with Cavity Ring Down Spectroscopy (Picarro, Inc., Santa Clara, CA, USA) in order to determine source water variability of the borehole fluids.

### Molecular Microbial Analysis

DNA from filters used to collect biomass from borehole fluids and soil samples were extracted with the ZymoBIOMICS DNA Miniprep Kit (Zymo Research, Irvine, CA, USA) and was used according to the manufacturer’s instructions. The concentration of raw DNA extract was quantified with the Qubit dsDNA high sensitivity assay (Thermo Fisher Scientific, Waltham, MA, USA). PCR amplification of the V4-V5 region of 16S rRNA gene was performed with the 515-Y-M13 forward (5’-**GTA AAA CGA CGG CCA GT**C CGT GYC AGC MGC CGC GTT AA-3’) and 926R reverse (5’-CCG YCA ATT YMT TTR AGT TT-3’) primers. The reverse primer was illustrated by Parada et al [58], and the forward primer contains the M13 forward primer sequence (outlined in bold) to aid in an additional PCR amplification for barcoding attachment (for subsequent DNA sequencing) along with the 16S rRNA gene specific sequence (the remaining sequence not in bold), as performed by [61, 62]. A second PCR amplification step was then included for the addition of barcodes to the M13 region of the forward primer as adapted from [63]. Initial PCR was performed on a Techne-TC 5000 thermocycler with the following parameters: initial denaturation at 94°C for 2 minutes; followed by 30 cycles of 94°C for 45 seconds, 50°C for 45 seconds, and 68°C for 1:30 minutes; ending with a final extension at 68°C for 5 minutes and a hold at 4°C until removal from the thermocycler. Barcoding of sequences was performed on a limited 6-cycle PCR on the purified initial PCR product. PCR product cleanup was performed with a 0.8x concentration of KAPA Pure Beads (KAPA Biosystems, Indianapolis, IN USA) according to the manufacturer’s instructions. Sequencing was performed on an Illumina MiSeq at either the Duke Center for Genomic or the Computational Biology (Raleigh, North Carolina) or the University of Colorado, Denver, Anschutz Medical Campus. Reads were processed using a modified pipeline illustrated by [64] for merging paired-end reads, quality filtering, constructing a sequence table, and removing chimeric sequences. Taxonomy was assigned suing the SILVA SSU database training file (version 138). ASV tables, phylogenetic trees, and sample metadata were collected into a phyloseq object for further downstream analysis [65].

### Statistical Analysis

All microbial community and geochemical multivariate analyses were conducted and produced in R. To assess microbial communities of the boreholes, all sample reads were rarified to an even depth of 5,761 reads per sample to remove as few reads as possible while maintaining enough diversity amongst samples—threshold determined by rarefaction curves (**Figure S4**). The samples were also ordinated though Principal Coordinate Analysis (PCoA) with a weighted Unifrac distance measure in order to consider phylogenetic relatedness as well as relative abundances. All figures were generated and visualized with the “ggplot2” or “phlyosmith” packages [66–68].

### Data Accessibility

The raw, unmerged 16S amplicon sequences as forward and reverse read files are publicly available on Figshare at 10.6084/m9.figshare.14740791. The code used for this study and additional metadata are available from our Github page: https://github.com/pthieringer/Edgar-Mine.

## Acknowledgments

We thank Matt Schreiner, Lee Fronapfel, and Clinton Dattel for insightful collaboration with safety training and experimental design at the Edgar Experimental Mine. We are grateful to Katherine Dawson at Rutgers University for guidance and analysis of water isotope data.

This work is supported by National Science Foundation Graduate Research Fellowships (Grant No. 1646713): Fellow I.D.s #2018254777 (P.H.T.), and #2019258966 (A.S.H.). The funders had no role in study design, data collection and analysis, decision to submit for publication, or preparation of the manuscript.

We declare that the research herein as conducted in the absence of any commercial or financial relationships that could act as a potential conflict of interest.

---

**Figure S1.**
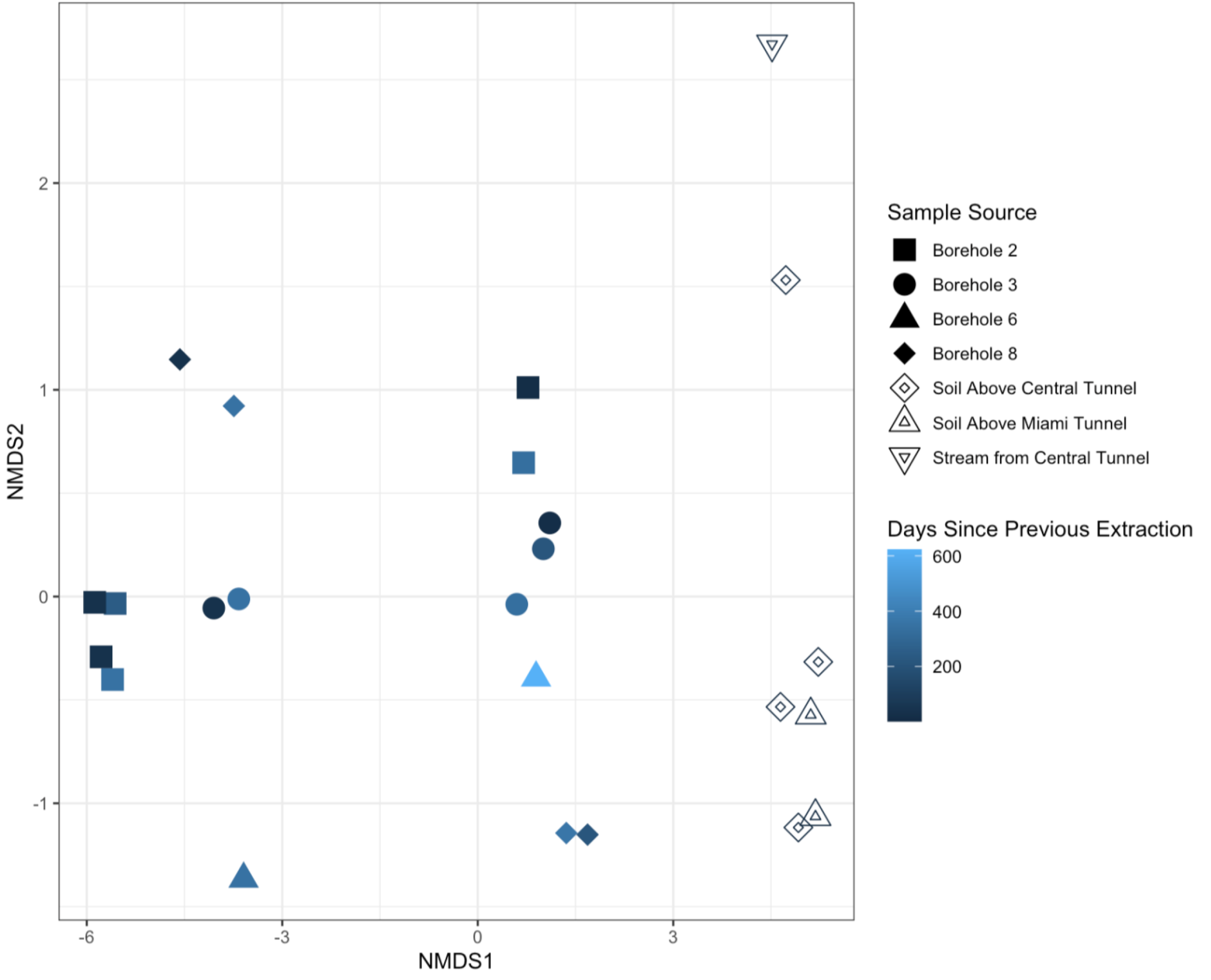
NMDS Plot of time since previous extraction for each sample. Clustering does not appear to show any distinct trends or associations. Soil samples are indicated as such and do not include a time variable due to only a single collection period. An Adonis test demonstrates that the time since the previous extraction for each sample is not significant (R^2^ = 0.073, p = 0.05).

**Figure S2.**
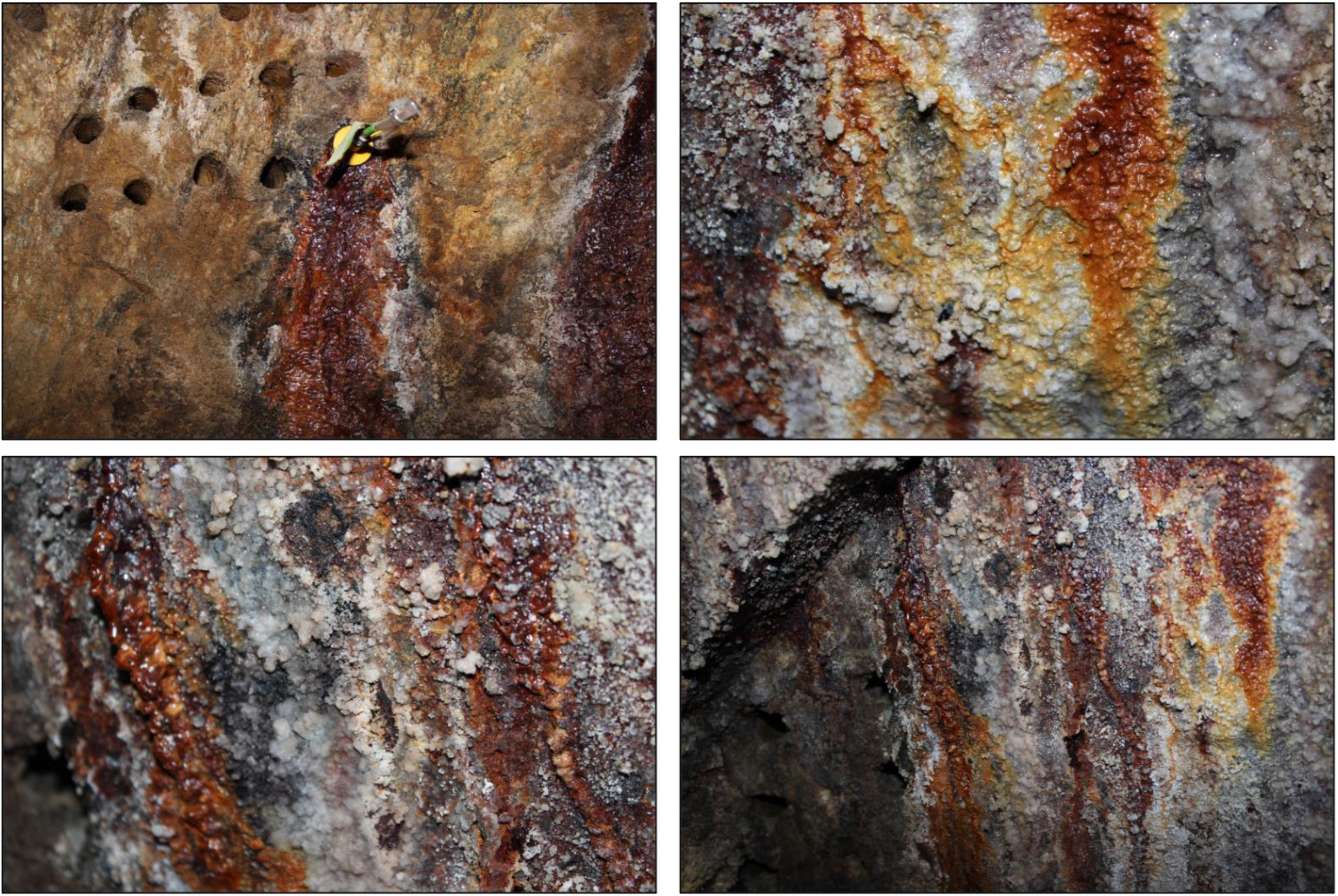
Potential biofilms collecting along the rock faces along the walls of the mine from leaking boreholes that are not packed. Discoloration from white to dark brown can be observed with crystal-like precipitates forming along the leakage path.

**Figure S3.**
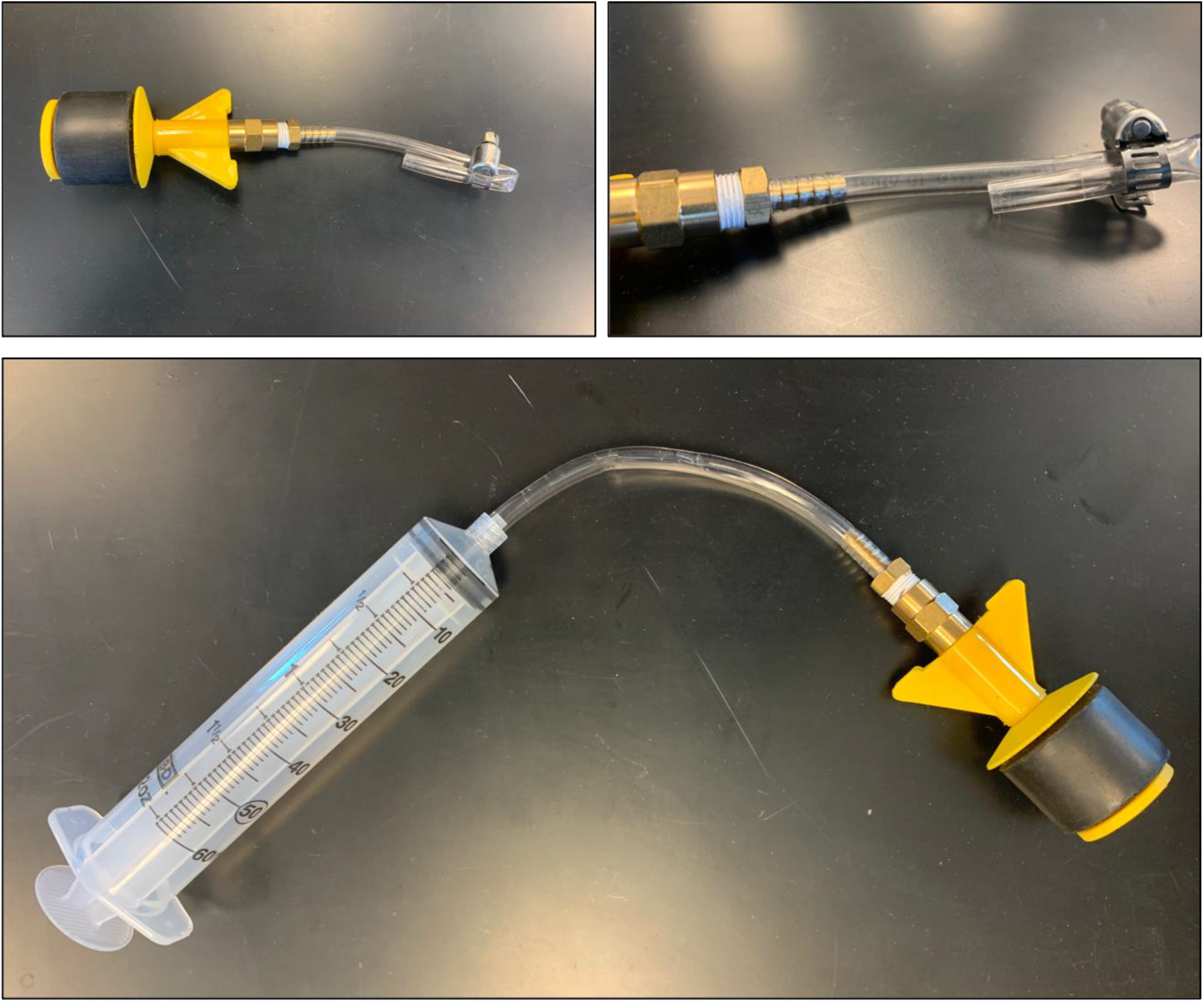
Packer contraption used to seal boreholes with leaking fluid. The black rubber end is inserted into the borehole to trap fluid, while the plastic housing is crimped to prevent any fluid from flowing out. The housing can then be attached to a syringe in order to pull fluid out of the borehole for biological or geochemical sampling purposes.

**Figure S4.**
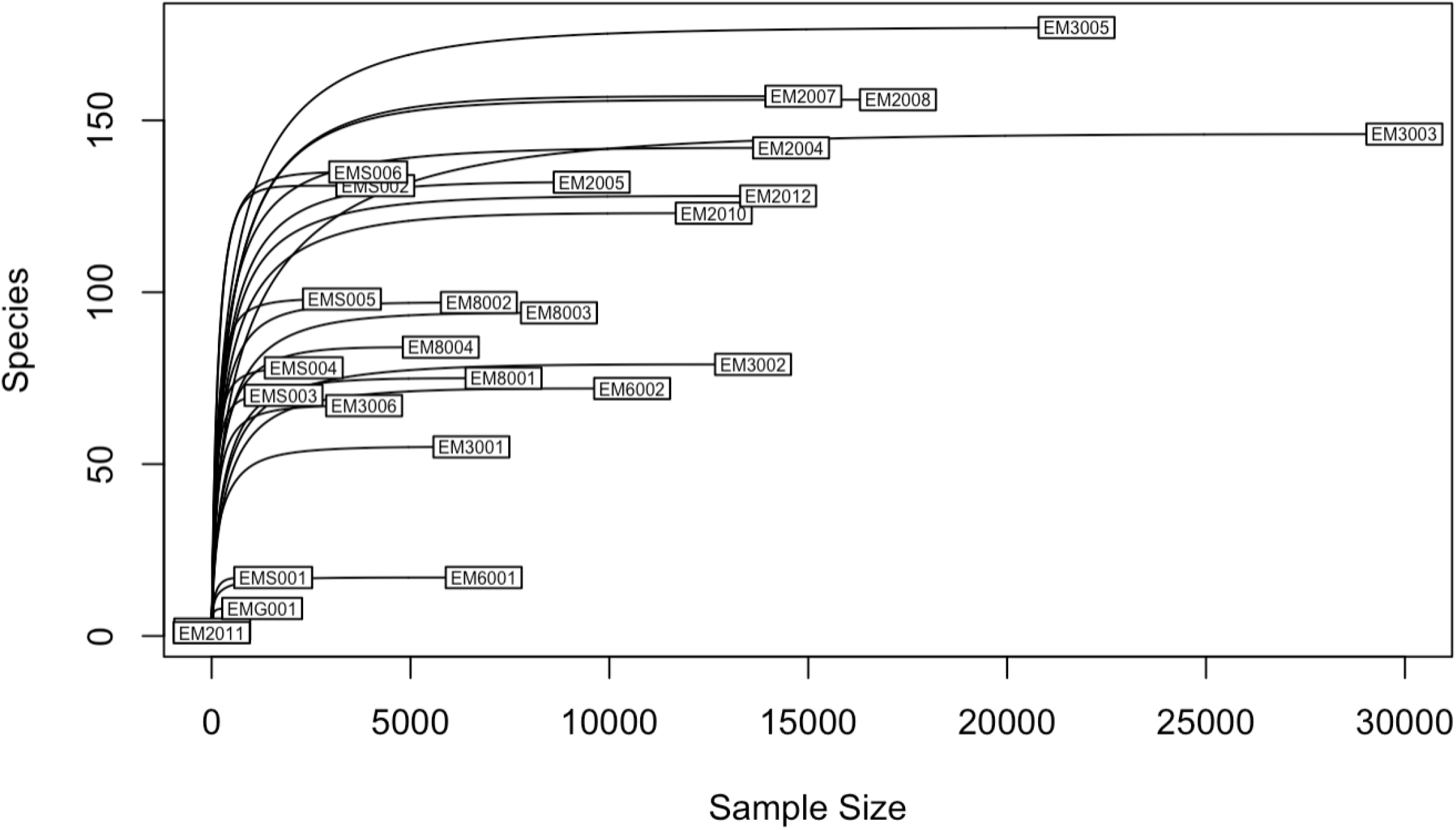
Rarefaction curves of samples collected from borehole and soil samples. Each curve is representative of an individual sample with the diversity mean species richness plotted on the y-axis and number of sequences on the x-axis. A rarefaction value of 5,761 sequences for the dataset used in this study to observe all borehole fluid samples, a rarefaction value of 1,270 was used to observe borehole fluids and other samples for ordination.

## References

[1] Gold, Thomas. 1992. “The Deep, Hot Biosphere | PNAS.” March 13, 1992. https://www.pnas.org/content/89/13/6045.

[2] Michalski, Joseph R., Tullis C. Onstott, Stephen J. Mojzsis, John Mustard, Queenie H. S. Chan, Paul B. Niles, and Sarah Stewart Johnson. 2018. “The Martian Subsurface as a Potential Window into the Origin of Life.” Nature Geoscience 11 (1): 21–26. https://doi.org/10.1038/s41561-017-0015-2

[3] Spear, John R, Jeffrey J Walker, Thomas M McCollom, and Norman R Pace. 2005. “Hydrogen and Bioenergetics in the Yellowstone Geothermal Ecosystem,” PNAS 102 (7) 2555–2560; https://doi.org/10.1073/pnas.0409574102

[4] Lynch, Kennda L., Briony H. Horgan, Junko Munakata-Marr, Jennifer Hanley, Robin J. Schneider, Kevin A. Rey, John R. Spear, W. Andrew Jackson, and Scott M. Ritter. 2015. “Near-Infrared Spectroscopy of Lacustrine Sediments in the Great Salt Lake Desert: An Analog Study for Martian Paleolake Basins: Mars Analog Paleolake Spectroscopy.” Journal of Geophysical Research: Planets 120 (3): 599–623. https://doi.org/10.1002/2014JE004707

[5] Kelley, Deborah S., Jeffrey A. Karson, Gretchen L. Früh-Green, Dana R. Yoerger, Timothy M. Shank, David A. Butterfield, John M. Hayes, et al. 2005. “A Serpentinite-Hosted Ecosystem: The Lost City Hydrothermal Field.” Science, New Series 307 (5714): 1428–34.

[6] Trembath-Reichert, Elizabeth, Yuki Morono, Akira Ijiri, Tatsuhiko Hoshino, Katherine S. Dawson, Fumio Inagaki, and Victoria J. Orphan. 2017. “Methyl-Compound Use and Slow Growth Characterize Microbial Life in 2-Km-Deep Subseafloor Coal and Shale Beds.” Proceedings of the National Academy of Sciences 114 (44): E9206–15. https://doi.org/10.1073/pnas.1707525114.

[7] Colman, Daniel R., Saroj Poudel, Blake W. Stamps, Eric S. Boyd, and John R. Spear. 2017. “The Deep, Hot Biosphere: Twenty-Five Years of Retrospection.” Proceedings of the National Academy of Sciences 114 (27): 6895–6903. https://doi.org/10.1073/pnas.1701266114

[8] Holland, G., B. Sherwood Lollar, L. Li, G. Lacrampe-Couloume, G. F. Slater, and C. J. Ballentine. 2013. “Deep Fracture Fluids Isolated in the Crust since the Precambrian Era.” Nature 497 (7449): 357–60. https://doi.org/10.1038/nature12127.

[9] Brazelton, William J., Christopher N. Thornton, Alex Hyer, Katrina I. Twing, August A. Longino, Susan Q. Lang, Marvin D. Lilley, Gretchen L. Früh-Green, and Matthew O. Schrenk. 2017. “Metagenomic Identification of Active Methanogens and Methanotrophs in Serpentinite Springs of the Voltri Massif, Italy.” PeerJ 5 (January): e2945. https://doi.org/10.7717/peerj.2945.

[10] Rempfert, Kaitlin R., Hannah M. Miller, Nicolas Bompard, Daniel Nothaft, Juerg M. Matter, Peter Kelemen, Noah Fierer, and Alexis S. Templeton. 2017. “Geological and Geochemical Controls on Subsurface Microbial Life in the Samail Ophiolite, Oman.” Frontiers in Microbiology 8 (February). https://doi.org/10.3389/fmicb.2017.00056.

[11] Kraus, Emily A., Daniel Nothaft, Blake W. Stamps, Kaitlin R. Rempfert, Eric T. Ellison, Juerg M. Matter, Alexis S. Templeton, Eric S. Boyd, and John R. Spear. 2020. “Molecular Evidence for an Active Microbial Methane Cycle in Subsurface Serpentinite-Hosted Groundwaters in the Samail Ophiolite, Oman.” Edited by Haruyuki Atomi. Applied and Environmental Microbiology 87 (2): e02068–20, /aem/87/2/AEM.02068-20.atom. https://doi.org/10.1128/AEM.02068-20.

[12] McMahon, Sean, and John Parnell. 2014. “Weighing the Deep Continental Biosphere.” FEMS Microbiology Ecology 87 (1): 113–20. https://doi.org/10.1111/1574-6941.12196.

[13] Schulte, Mitch, David Blake, Tori Hoehler, and Thomas McCollom. 2006. “Serpentinization and Its Implications for Life on the Early Earth and Mars.” Astrobiology 6 (2): 364–76. https://doi.org/10.1089/ast.2006.6.364

[14] Barnes, Ivan and O’Neil, James. 1969. “The Relationship between Fluids in Some Fresh Alpine-Type Ultramafics and Possible Modern Serpentinization, Western United States.” Geological Society of America Bulltetin 80 (October): 19471960.

[15] Osburn, Magdalena R., Douglas E. LaRowe, Lily M. Momper, and Jan P. Amend. 2014. “Chemolithotrophy in the Continental Deep Subsurface: Sanford Underground Research Facility (SURF), USA.” Frontiers in Microbiology 5 (November). https://doi.org/10.3389/fmicb.2014.00610

[16] Baker, Brett J, and Jillian F Banfield. 2003. “Microbial Communities in Acid Mine Drainage.” FEMS Microbiology Ecology 44 (2): 139–52. https://doi.org/10.1016/S0168-6496(03)00028-X

[17] Onstott, T. C., L.-H. Lin, M. Davidson, B. Mislowack, M. Borcsik, J. Hall, G. Slater, et al. 2006. “The Origin and Age of Biogeochemical Trends in Deep Fracture Water of the Witwatersrand Basin, South Africa.” Geomicrobiology Journal 23 (6): 369–414. https://doi.org/10.1080/01490450600875688

[18] Schrenk, M. O., W. J. Brazelton, and S. Q. Lang. 2013. “Serpentinization, Carbon, and Deep Life.” Reviews in Mineralogy and Geochemistry 75 (1): 575–606. https://doi.org/10.2138/rmg.2013.75.18

[19] Lollar, Garnet S., Oliver Warr, Jon Telling, Magdalena R. Osburn, and Barbara Sherwood Lollar. 2019. “‘Follow the Water’: Hydrogeochemical Constraints on Microbial Investigations 2.4 Km Below Surface at the Kidd Creek Deep Fluid and Deep Life Observatory.” Geomicrobiology Journal 36 (10): 859–72. https://doi.org/10.1080/01490451.2019.1641770

[20] Onstott, T. C., D. P. Moser, S. M. Pfiffner, J. K. Fredrickson, F. J. Brockman, T. J. Phelps, D. C. White, et al. 2003. “Indigenous and Contaminant Microbes in Ultradeep Mines.” Environmental Microbiology 5 (11): 1168–91. https://doi.org/10.1046/j.1462-2920.2003.00512.x

[21] Kendall, Carol, and Tyler B. Coplen. 2001. “Distribution of Oxygen-18 and Deuterium in River Waters across the United States.” Hydrological Processes 15 (7): 1363–93. https://doi.org/10.1002/hyp.217

[22] Hallbeck, Lotta, and Karsten Pedersen. 2014. “The Family Gallionellaceae.” In The Prokaryotes, edited by Eugene Rosenberg, Edward F. DeLong, Stephen Lory, Erko Stackebrandt, and Fabiano Thompson, 853–58. Berlin, Heidelberg: Springer Berlin Heidelberg. https://doi.org/10.1007/978-3-642-30197-1_398

[23] Kadnikov, V. V., D. A. Ivasenko, A. V. Beletskii, A. V. Mardanov, E. V. Danilova, N. V. Pimenov, O. V. Karnachuk, and N. V. Ravin. 2016. “A Novel Uncultured Bacterium of the Family Gallionellaceae: Description and Genome Reconstruction Based on Metagenomic Analysis of Microbial Community in Acid Mine Drainage.” Microbiology

[24] Kojima, Hisaya, Arisa Shinohara, and Manabu Fukui. 2015. “Sulfurifustis Variabilis Gen. Nov., Sp. Nov., a Sulfur Oxidizer Isolated from a Lake, and Proposal of Acidiferrobacteraceae Fam. Nov. and Acidiferrobacterales Ord. Nov.” International Journal of Systematic and Evolutionary Microbiology 65 (Pt_10): 3709–13. https://doi.org/10.1099/ijsem.0.000479

[25] Alazard, Didier, Manon Joseph, Fabienne Battaglia-Brunet, Jean-Luc Cayol, and Bernard Ollivier. 2010. “Desulfosporosinus Acidiphilus Sp. Nov.: A Moderately Acidophilic Sulfate-Reducing Bacterium Isolated from Acid Mining Drainage Sediments: New Taxa: Firmicutes (Class Clostridia, Order Clostridiales, Family Peptococcaceae).” Extremophiles 14 (3): 305–12. https://doi.org/10.1007/s00792-010-0309-4.

[26] Glöckner, Jana, Michael Kube, Pravin Malla Shrestha, Marc Weber, Frank Oliver Glöckner, Richard Reinhardt, and Werner Liesack. 2010. “Phylogenetic Diversity and Metagenomics of Candidate Division OP3: Candidate Division OP3.” Environmental Microbiology 12 (5): 1218–29. https://doi.org/10.1111/j.1462-2920.2010.02164.x.

[27] Egli, Thomas W., and Georg Auling. 2005. “Chelatococcus Auling, Busse, Egli, El-Banna and Stackebrandt 1993b, 624VP (Effective Publication: Auling, Busse, Egli, El-Banna and Stackebrandt 1993a, 109).” In Bergey’s Manual of Systematic Bacteriology, edited by Don J. Brenner, Noel R. Krieg, George M. Garrity, James T. Staley, David R. Boone, Paul Vos, Michael Goodfellow, Fred A. Rainey, and Karl-Heinz Schleifer, 433–37. New York: Springer-Verlag. https://doi.org/10.1007/0-387-29298-5_105

[28] Birtles, R.J., and D. Raoult. 1998. “The Genera *Afipia* and *Bartonella*.” In Contributions to Microbiology, edited by A. Schmidt, 1:1–31. Basel: KARGER. https://doi.org/10.1159/000060452

[29] Kim, Yeon-Ju, Ngoc-Lan Nguyen, Hang-Yeon Weon, and Deok-Chun Yang. n.d. “Sediminibacterium Ginsengisoli Sp. Nov., Isolated from Soil of a Ginseng Field, and Emended Descriptions of the Genus Sediminibacterium and of Sediminibacterium Salmoneum.” International Journal of Systematic and Evolutionary Microbiology, 8.

[30] Li, Liqiong, Hongliang Liu, Zunji Shi, and Gejiao Wang. 2013. “Sphingobium Cupriresistens Sp. Nov., a Copper-Resistant Bacterium Isolated from Copper Mine Soil, and Emended Description of the Genus Sphingobium.” International Journal of Systematic and Evolutionary Microbiology 63 (Pt_2): 604–9. https://doi.org/10.1099/ijs.0.040865-0.

[31] Ormeño-Orrillo, Ernesto, and Esperanza Martínez-Romero. 2019. “A Genomotaxonomy View of the Bradyrhizobium Genus.” Frontiers in Microbiology 10 (June): 1334. https://doi.org/10.3389/fmicb.2019.01334

[32] Whitman, William B, Fred Rainey, Peter Kämpfer, Martha Trujillo, Jonsik Chun, Paul DeVos, Brian Hedlund, and Svetlana Dedysh, eds. 2015. Candidatus Nitrosotenuis. 1st ed. Wiley. https://doi.org/10.1002/9781118960608

[33] Xing, Tingting, Tandong Yao, Yongqin Liu, Ninglian Wang, Bainqing Xu, Liang Shen, Zhengquan Gu, Bixi Gu, Hongcan Liu, and Yuguang Zhou. 2016. “Polaromonas Eurypsychrophila Sp. Nov., Isolated from an Ice Core.” International Journal of Systematic and Evolutionary Microbiology 66 (7): 2497–2501. https://doi.org/10.1099/ijsem.0.001079

[34] Gawor, Jan, Jakub Grzesiak, Joanna Sasin-Kurowska, Piotr Borsuk, Robert Gromadka, Dorota Górniak, Aleksander Świątecki, Tamara Aleksandrzak-Piekarczyk, and Marek K. Zdanowski. 2016. “Evidence of Adaptation, Niche Separation and Microevolution within the Genus Polaromonas on Arctic and Antarctic Glacial Surfaces.” Extremophiles 20 (4): 403–13. https://doi.org/10.1007/s00792-016-0831-0.

[35] Lozupone, Catherine, and Rob Knight. 2005. “UniFrac: A New Phylogenetic Method for Comparing Microbial Communities.” Applied and Environmental Microbiology 71 (12): 8228–35. https://doi.org/10.1128/AEM.71.12.8228-8235.2005

[36] Warr, Oliver, Thomas Giunta, Tullis C. Onstott, Thomas L. Kieft, Rachel L. Harris, Devan M. Nisson, and Barbara Sherwood Lollar. 2021. “The Role of Low-Temperature 18O Exchange in the Isotopic Evolution of Deep Subsurface Fluids.” Chemical Geology 561 (February): 120027. https://doi.org/10.1016/j.chemgeo.2020.120027

[37] Lau, Maggie C. Y., Thomas L. Kieft, Olukayode Kuloyo, Borja Linage-Alvarez, Esta van Heerden, Melody R. Lindsay, Cara Magnabosco, et al. 2016. “An Oligotrophic Deep-Subsurface Community Dependent on Syntrophy Is Dominated by Sulfur-Driven Autotrophic Denitrifiers.” Proceedings of the National Academy of Sciences 113 (49): E7927–36. https://doi.org/10.1073/pnas.1612244113

[38] Bell, Emma, Tiina Lamminmäki, Johannes Alneberg, Anders F. Andersson, Chen Qian, Weili Xiong, Robert L. Hettich, Manon Frutschi, and Rizlan Bernier-Latmani. 2020. “Active Sulfur Cycling in the Terrestrial Deep Subsurface.” The ISME Journal 14 (5): 1260–72. https://doi.org/10.1038/s41396-020-0602-x.

[39] Stein, Otto R., Deborah J. Borden-Stewart, Paul B. Hook, and Warren L. Jones. 2007. “Seasonal Influence on Sulfate Reduction and Zinc Sequestration in Subsurface Treatment Wetlands.” Water Research 41 (15): 3440–48. https://doi.org/10.1016/j.watres.2007.04.023

[40] Ghosh, Shreya, and Alok Prasad Das. 2018. “Metagenomic Insights into the Microbial Diversity in Manganese-Contaminated Mine Tailings and Their Role in Biogeochemical Cycling of Manganese.” Scientific Reports 8 (1): 8257. https://doi.org/10.1038/s41598-018-26311-w.

[41] Bal, Bhubaneswari, Shreya Ghosh, and Alok Prasad Das. 2019. “Microbial Recovery and Recycling of Manganese Waste and Their Future Application: A Review.” Geomicrobiology Journal 36 (1): 85–96

[42] Sahl, Jason W., Raleigh Schmidt, Elizabeth D. Swanner, Kevin W. Mandernack, Alexis S. Templeton, Thomas L. Kieft, Richard L. Smith, et al. 2008. “Subsurface Microbial Diversity in Deep-Granitic-Fracture Water in Colorado.” Applied and Environmental Microbiology 74 (1): 143–52. https://doi.org/10.1128/AEM.01133-07

[43] Lohmann, Patrick, Simon Benk, Gerd Gleixner, Karin Potthast, Beate Michalzik, Nico Jehmlich, and Martin von Bergen. 2020. “Seasonal Patterns of Dominant Microbes Involved in Central Nutrient Cycles in the Subsurface.” Microorganisms 8 (11): 1694. https://doi.org/10.3390/microorganisms8111694.

[44] Blume, E., M. Bischoff, J.M. Reichert, T. Moorman, A. Konopka, and R.F. Turco. 2002. “Surface and Subsurface Microbial Biomass, Community Structure and Metabolic Activity as a Function of Soil Depth and Season.” Applied Soil Ecology 20 (3): 171–81. https://doi.org/10.1016/S0929-1393(02)00025-2.

[45] Honeyman, Alexander S., Maria L. Day, and John R. Spear. 2018. “Regional Fresh Snowfall Microbiology and Chemistry Are Driven by Geography in Storm-Tracked Events, Colorado, USA.” PeerJ 6 (November): e5961. https://doi.org/10.7717/peerj.5961

[46] Fredrickson, James K., and David L. Balkwill. 2006. “Geomicrobial Processes and Biodiversity in the Deep Terrestrial Subsurface.” Geomicrobiology Journal 23 (6): 345–56. https://doi.org/10.1080/01490450600875571

[47] Munson-McGee, Jacob H., Erin K. Field, Mary Bateson, Colleen Rooney, Ramunas Stepanauskas, and Mark J. Young. 2015. “Nanoarchaeota, Their Sulfolobales Host, and Nanoarchaeota Virus Distribution across Yellowstone National Park Hot Springs.” Edited by K. E. Wommack. Applied and Environmental Microbiology 81 (22): 7860–68. https://doi.org/10.1128/AEM.01539-15.

[48] Huber, Harald, Michael J Hohn, Karl O Stetter, and Reinhard Rachel. 2003. “The Phylum Nanoarchaeota: Present Knowledge and Future Perspectives of a Unique Form of Life.” Research in Microbiology 154 (3): 165–71. https://doi.org/10.1016/S0923-2508(03)00035-4.

[49] Wurch, Louie, Richard J. Giannone, Bernard S. Belisle, Carolyn Swift, Sagar Utturkar, Robert L. Hettich, Anna-Louise Reysenbach, and Mircea Podar. 2016. “Genomics-Informed Isolation and Characterization of a Symbiotic Nanoarchaeota System from a Terrestrial Geothermal Environment.” Nature Communications 7 (1): 12115. https://doi.org/10.1038/ncomms12115

[50] Sajjad, Wasim, Guodong Zheng, Ghufranud Din, Xiangxian Ma, Muhammad Rafiq, and Wang Xu. 2019. “Metals Extraction from Sulfide Ores with Microorganisms: The Bioleaching Technology and Recent Developments.” Transactions of the Indian Institute of Metals 72 (3): 559–79. https://doi.org/10.1007/s12666-018-1516-4.

[51] Bach, Wolfgang, and Katrina J Edwards. 2003. “Iron and Sulfide Oxidation within the Basaltic Ocean Crust: Implications for Chemolithoautotrophic Microbial Biomass Production.” Geochimica et Cosmochimica Acta 67 (20): 3871–87. https://doi.org/10.1016/S0016-7037(03)00304-1.

[52] Kitzinger, Katharina, Hanna Koch, Sebastian Lücker, Christopher J. Sedlacek, Craig Herbold, Jasmin Schwarz, Anne Daebeler, et al. 2018. “Characterization of the First ‘ *Candidatus* Nitrotoga’ Isolate Reveals Metabolic Versatility and Separate Evolution of Widespread Nitrite-Oxidizing Bacteria.” Edited by Douglas G. Capone. MBio 9 (4): e01186–18, /mbio/9/4/mBio.01186-18.atom. https://doi.org/10.1128/mBio.01186-18

[53] Probst, Alexander J., Giovanni Birarda, Hoi-Ying N. Holman, Todd Z. DeSantis, Gerhard Wanner, Gary L. Andersen, Alexandra K. Perras, et al. 2014. “Coupling Genetic and Chemical Microbiome Profiling Reveals Heterogeneity of Archaeome and Bacteriome in Subsurface Biofilms That Are Dominated by the Same Archaeal Species.” Edited by Gabriele Berg. PLoS ONE 9 (6): e99801. https://doi.org/10.1371/journal.pone.0099801

[54] Jones, D. S., D. J. Tobler, I. Schaperdoth, M. Mainiero, and J. L. Macalady. 2010. “Community Structure of Subsurface Biofilms in the Thermal Sulfidic Caves of Acquasanta Terme, Italy.” Applied and Environmental Microbiology 76 (17): 5902–10. https://doi.org/10.1128/AEM.00647-10

[55] Coombs, P., D. Wagner, K. Bateman, H. Harrison, A.E. Milodowski, D. Noy, and J.M. West. 2010. “The Role of Biofilms in Subsurface Transport Processes.” Quarterly Journal of Engineering Geology and Hydrogeology 43 (2): 131–39. https://doi.org/10.1144/1470-9236/08-029.

[56] Casar, Caitlin P., Brittany R. Kruger, Theodore M. Flynn, Andrew L. Masterson, Lily M. Momper, and Magdalena R. Osburn. 2020. “Mineral-hosted Biofilm Communities in the Continental Deep Subsurface, Deep Mine Microbial Observatory, SD, USA.” Geobiology 18 (4): 508–22. https://doi.org/10.1111/gbi.12391

[57] Lee, Jaeheon, Sevket Acar, Denise L. Doerr, and James A. Brierley. 2011. “Comparative Bioleaching and Mineralogy of Composited Sulfide Ores Containing Enargite, Covellite and Chalcocite by Mesophilic and Thermophilic Microorganisms.” Hydrometallurgy 105 (3–4): 213–21. https://doi.org/10.1016/j.hydromet.2010.10.001

[58] Giaveno, A., L. Lavalle, P. Chiacchiarini, and E. Donati. 2007. “Bioleaching of Zinc from Low-Grade Complex Sulfide Ores in an Airlift by Isolated Leptospirillum Ferrooxidans.” Hydrometallurgy 89 (1–2): 117–26. https://doi.org/10.1016/j.hydromet.2007.07.002

[59] Bosecker, Klaus. 1997. “Bioleaching: Metal Solubilization by Microorganisms.” FEMS Microbiology Reviews 20 (3–4): 591–604. https://doi.org/10.1111/j.1574-6976.1997.tb00340.x

[60] Mosch, David K., Petr, Vilem. 2013. “Underground Opening and Support Facilities of the Edgar Experimental Mine Idaho Springs, Colorado.” Colorado School of Mines. https://www.mines.edu/mining/wp-content/uploads/sites/28/2017/08/edgar-mine-information-2013.pdf

[61] Parada, Alma E., David M. Needham, and Jed A. Fuhrman. 2016. “Every Base Matters: Assessing Small Subunit RRNA Primers for Marine Microbiomes with Mock Communities, Time Series and Global Field Samples: Primers for Marine Microbiome Studies.” Environmental Microbiology 18 (5): 1403–14. https://doi.org/10.1111/1462-2920.13023

[62] Stamps, Blake W., Christopher N. Lyles, Joseph M. Suflita, Jason R. Masoner, Isabelle M. Cozzarelli, Dana W. Kolpin, and Bradley S. Stevenson. 2016. “Municipal Solid Waste Landfills Harbor Distinct Microbiomes.” Frontiers in Microbiology 7. https://doi.org/10.3389/fmicb.2016.00534

[63] Caporaso, J Gregory, Christian L Lauber, William A Walters, Donna Berg-Lyons, James Huntley, Noah Fierer, Sarah M Owens, et al. 2012. “Ultra-High-Throughput Microbial Community Analysis on the Illumina HiSeq and MiSeq Platforms.” The ISME Journal 6 (8): 1621–24. https://doi.org/10.1038/ismej.2012.8

[64] Callahan, Benjamin J, Paul J McMurdie, Michael J Rosen, Andrew W Han, Amy Jo A Johnson, and Susan P Holmes. 2016. “DADA2: High-Resolution Sample Inference from Illumina Amplicon Data.” Nature Methods 13 (7): 581–83. https://doi.org/10.1038/nmeth.3869.

[65] McMurdie, Paul J., and Susan Holmes. 2013. “Phyloseq: An R Package for Reproducible Interactive Analysis and Graphics of Microbiome Census Data.” Edited by Michael Watson. PLoS ONE 8 (4): e61217. https://doi.org/10.1371/journal.pone.0061217

[66] Wickham, Hadley. 2011. “Ggplot2.” Wiley Interdisciplinary Reviews: Computational Statistics 3 (2): 180–85. https://doi.org/10.1002/wics.147

[67] Smith, (2019). phylosmith: an R-package for reproducible and efficient microbiome analysis with phyloseq-objects. Journal of Open Source Software, 4(38), 1442, https://doi.org/10.21105/joss.01442

[68] R Development Core Team (2015). R: A Language and Environment for Statistical Computing. Vienna: R Foundation for tatistical Computing. Available online at: http://www.R-project.org/

